# Deoxyribonucleotide dephosphorylation by VENOSA4 supports organellar genome replication in plants

**DOI:** 10.64898/2026.09.08.750075

**Authors:** Lisa Fischer, Henryk Straube, Nele Passon, Hildegard Thölke, Claus-Peter Witte, Marco Herde

## Abstract

The replication of the three genomes in plant cells during germination is a complex process, which requires a high degree of coordination between the genome-containing compartments: the nucleus, mitochondria, and chloroplasts. The first committed step for the *de novo* synthesis of deoxynucleoside triphosphates (dNTPs), the building blocks for DNA replication, occurs exclusively in the cytosol. A major unresolved question is how an adequate supply of dNTPs for the organelles is achieved. Here, we show that VENOSA4 (VEN4), a dNTP triphosphohydrolase, is critical for this process in *Arabidopsis thaliana*. Using isotope feeding combined with mass spectrometry analysis, we demonstrate that VEN4 converts most *de novo*-synthesized dNTPs to deoxynucleosides (dNs). Preventing this conversion by *VEN4* mutation strongly diminishes both cpDNA and mtDNA amounts, but these can be partially rescued by the application of exogenous dNs. Hence, the dNTP catabolic activity of VEN4 ensures that sufficient DNA precursors in form of dNs reach the chloroplasts and the mitochondria to be salvaged there to dNTPs for DNA synthesis. This seemingly counterintuitive coupling of *de novo* synthesis, dNTP hydrolysis, and salvage may provide a mechanism to control DNA precursor allocation between cellular compartments through selective transport and metabolic trapping.

## Introduction

In plant cells, DNA replication occurs in the nucleus, the chloroplasts, and the mitochondria (Draper and Hays 2000). Although the two organelles possess their own genome and replication machinery (Morley et al. 2019), the synthesis of deoxyribonucleotides (dNTPs), the essential substrates for DNA replication and repair, is located primarily in the cytosol (Wang and Liu 2006; Witte and Herde 2020). This spatial separation requires that cytosolic dNTP production must be synchronized with the replication activity of all three genomes (Draper and Hays 2000; Law et al. 2012). During germination and seedling establishment (GSE), when metabolic activity is rapidly increasing and organelle biogenesis is prominent, it is particularly important to coordinate the dNTP supply (Draper and Hays 2000; Law et al. 2012; Niehaus et al. 2022). How cytosolic dNTP pools are distributed, balanced, and supplied to organelles during GSE remains poorly understood (Niehaus et al. 2022; Witte and Herde 2020).

Plants obtain precursors for DNA synthesis and repair either from *de novo* dNTP synthesis or from salvage pathways (Witte and Herde 2020). The first committed step in *de novo* dNTP synthesis is exclusively located in the cytosol and catalyzed by ribonucleotide reductase (RNR) (Wang and Liu 2006). This multimeric enzyme complex reductively removes the 2’-hydroxyl moiety on the ribose of ribonucleoside diphosphates (rNDPs), resulting in the formation of deoxyribonucleoside diphosphates (dNDPs) (Wang and Liu 2006; Witte and Herde 2020). These are phosphorylated to form deoxyribonucleoside triphosphates (dNTPs) via nucleoside diphosphate kinases (NDPKs) (Dorion et al. 2006; Hammargren et al. 2007). NDPKs exhibit multi-substrate functionality, producing dATP, dCTP, dGTP, and dUTP in the cytosol, the chloroplasts, and the mitochondria (Bölter et al. 2007; Dorion et al. 2006; Hammargren et al. 2007; Sweetlove et al. 2001). While dATP, dCTP, and dGTP can be used directly for DNA synthesis, dUTP cannot; instead, dTTP must be formed. The generation of dTTP proceeds via dUMP formation, either through dephosphorylation of dUTP to dUMP by dUTP pyrophosphatase-like1 (DUT1) (Dubois et al. 2011) or through dephosphorylation of dUDP to dUMP by a putative unknown dUDP phosphatase (Niehaus et al. 2022; Witte and Herde 2020). A third option for dUMP production is the deamination of dCMP/dCTP to dUMP/dUTP by dCMP/dCTP deaminase (DCD) in mitochondria (Niu et al. 2017; Witte and Herde 2020; Xu et al. 2014). The methylation of dUMP to dTMP is then catalyzed either in the cytosol by dihydrofolate reductase-thymidylate synthase 1 (DHFR-TS1) or in mitochondria by dihydrofolate reductase-thymidylate synthase 2 (DHFR-TS2), but not in chloroplasts (Gorelova et al. 2017; Niehaus et al. 2022). Further phosphorylation of dTMP to dTTP is catalyzed by dTMP kinase (TMPK) and NDPK (Dorion et al. 2006; Hammargren et al. 2007; Ronceret et al. 2008). These phosphorylation reactions occur in the cytosol, in mitochondria, and in plastids (Bölter et al. 2007; Ronceret et al. 2008; Sweetlove et al. 2001). The RNR-dependent *de novo* synthesis of dNTPs is central for proper plant development. The *A. thaliana* RNR small-subunit mutant *tso-2* shows reduced dNTP concentrations, disrupted cell-cycle progression, and impaired DNA damage repair (Wang and Liu 2006). A null mutation of *SvSTL1*, a gene coding for a homolog of the RNR large subunit in *Setaria viridis*, resulted in impaired chloroplast development and reduced cpDNA copy number. The authors of that study proposed that a reduced cytosolic dNTP supply, which was not directly measured, may account for the decrease in cpDNA and that nuclear DNA replication is prioritized when cytosolic dNTP pools are limiting (Li et al. 2022).

Apart from *de-novo* synthesis, DNA building blocks can originate from salvage pathways. Salvage is the process by which nucleobases and nucleosides or deoxynucleosides (dNs) are reintroduced into the corresponding pools of nucleotide monophosphates (NMPs) or deoxynucleoside monophosphates (dNMPs) (Clausen et al. 2012; Witte and Herde 2020). It is often referred to as an energy-saving way to generate nucleotides by recycling building blocks from RNA/DNA repair (Girke et al. 2014; Le Ret et al. 2018; Witte and Herde 2020). dG, dC, dA and dU but not dT are salvaged to the corresponding dNMPs via phosphorylation by deoxynucleoside kinase (dNK) (Clausen et al. 2012). Phosphorylation of dT to dTMP is exclusively mediated by thymidine kinases (TKs) (Clausen et al., 2012; Le Ret et al., 2018). In Arabidopsis, two structurally similar TK isoforms with different subcellular localizations exist. TK1a is located in the cytosol and TK1b is present in mitochondria and chloroplasts (Clausen et al. 2012; Le Ret et al. 2018; Niehaus et al. 2022). A double null-mutant of *TK1a* and *TK1b* is seedling lethal, showing that dT salvage is essential for plant development. Why the TKs are so important is currently not understood (Clausen et al. 2012).

Clearly, dNTP *de novo* synthesis and salvage are essential for proper plant development. Nonetheless, they have often been considered independent processes that contribute separately to DNA synthesis. However, an earlier study from our group had already suggested that both processes are intertwined.

We showed that a null mutation of *TK1b* leads to a reduction in cpDNA by two-thirds during GSE. The source of the dT processed by TK1b was unclear but was not DNA repair, as previously assumed, because dT concentrations were unchanged in global genome repair mutants. Dephosphorylation of stored dTTP as a source of dT could also be excluded since dNTPs pools were shown to be very small in dry seeds (Niehaus et al. 2022). We speculated that *de novo* synthesis of dTTP could be the source of dT because during GSE dNTP and dT concentrations correlate - both increase with the onset of DNA replication. This suggests that cytosolic dNTP *de novo* synthesis supplies dNs to the organellar salvage pathway. Such a pathway is counterintuitive because dNTPs are first dephosphorylated and then re-phosphorylated, partially offsetting the energetic advantage of salvage. It would therefore require a dNTP-dephosphorylating enzyme that counteracts RNR-dependent dNTP synthesis. A possible candidate to catalyze this reaction is venosa4 (VEN4) in *A. thaliana*. VEN4 is a homologue of the mammalian sterile alpha motif and histidine-aspartate (HD) domain-containing protein 1 (SAMHD1), which hydrolyzes dNTPs into dNs and inorganic triphosphate (Franzolin et al. 2013; Goldstone et al. 2011). In mammals, this enzyme is primarily known as a regulator of dNTP homeostasis during the cell cycle (Franzolin et al. 2013). Outside S-phase, SAMHD1 maintains low cellular dNTP pools and thereby inhibits, for example, HIV replication (Goldstone et al. 2011; Lahouassa et al. 2012).

Null mutants of *VEN4* in Arabidopsis, have been associated with a chlorotic phenotype, reduced photosynthetic efficiency, and abnormal thylakoid membrane structures in chloroplasts (Sarmiento-Mañús et al. 2023; Xu et al. 2020; Yoshida et al. 2018). Because VEN4 is homologous to SAMHD1, its mutant phenotypes have been proposed to result from altered dNTP homeostasis (Sarmiento-Mañús et al. 2023; Wang et al. 2022; Xu et al. 2020). However, the effect of VEN4 loss on cellular dNTP pools has not been established. It is important to note that an intrinsic link between VEN4 or SAMHD1, dNTP *de novo* synthesis and salvage has not been clearly demonstrated in any organism so far.

In this study, our aim was to determine the origin of dT during GSE and to test whether VEN4 links cytosolic *de novo* dNTP synthesis to organellar dT salvage and, consequently, the dNTP supply to the organelles. To distinguish *de novo*-from salvage-derived thymidylates, we combined ^15^N-uridine labeling with liquid chromatography–tandem mass spectrometry (LC– MS/MS) and inhibition of dNTP and dTMP *de novo* synthesis. The concentrations of dNTPs and dNs as well as cpDNA and mtDNA abundance were analyzed in *VEN4* mutants to examine the contribution of VEN4 to dNTP metabolism and organellar DNA synthesis. Genetic interactions between *VEN4* and *TK1b* were analyzed to test whether VEN4-dependent dT formation contributes to organellar dT salvage. In addition, we investigated whether mitochondrial DHFR-TS2 and cytosolic TK1a can provide thymidylates when VEN4 function is impaired.

## Results

### 15N-uridine feeding labels pyrimidine pools and supports a *de novo* origin of thymidine

To estimate the flux through the two thymidylate-generating pathways—the *de novo* synthesis of dTTP and the salvage of dT to dTTP—we grew *A. thaliana* (Col-0) seeds for 48 hours on liquid ½ MS medium supplemented with stable isotope-labeled uridine (Uridine-1,3-^15^N_2_; hereafter ^15^N-uridine). Label incorporation into nucleotides was analyzed by LC–MS/MS. The label can enter the dNT pool via uridylates or cytidylates (Witte and Herde 2020) (Fig. 1A). Uridine is imported to plant cells by the equilibrative nucleoside transporter ENT3 (Chen et al. 2006), phosphorylated to UDP by uridine-cytidine kinases (UCKs) (Ohler et al. 2019; Witte and Herde 2020) and uridine monophosphate kinases (UMKs) (Rinne et al. 2024), and is then reduced to dUDP by RNR (Wang and Liu 2006) (Fig. 1A). Alternatively, UDP can be further phosphorylated to UTP by NDPKs (Dorion et al. 2006) and subsequently aminated by CTP synthase (CTPS) (Daumann et al. 2018; Hickl et al. 2021), thereby entering the cytidylate pool. The resulting CTP can then be dephosphorylated to CDP (Rinne et al. 2024; Witte and Herde 2020), which, in turn, can be reduced to dCDP by RNR (Wang and Liu 2006) (Fig. 1A).

**Figure 1.**
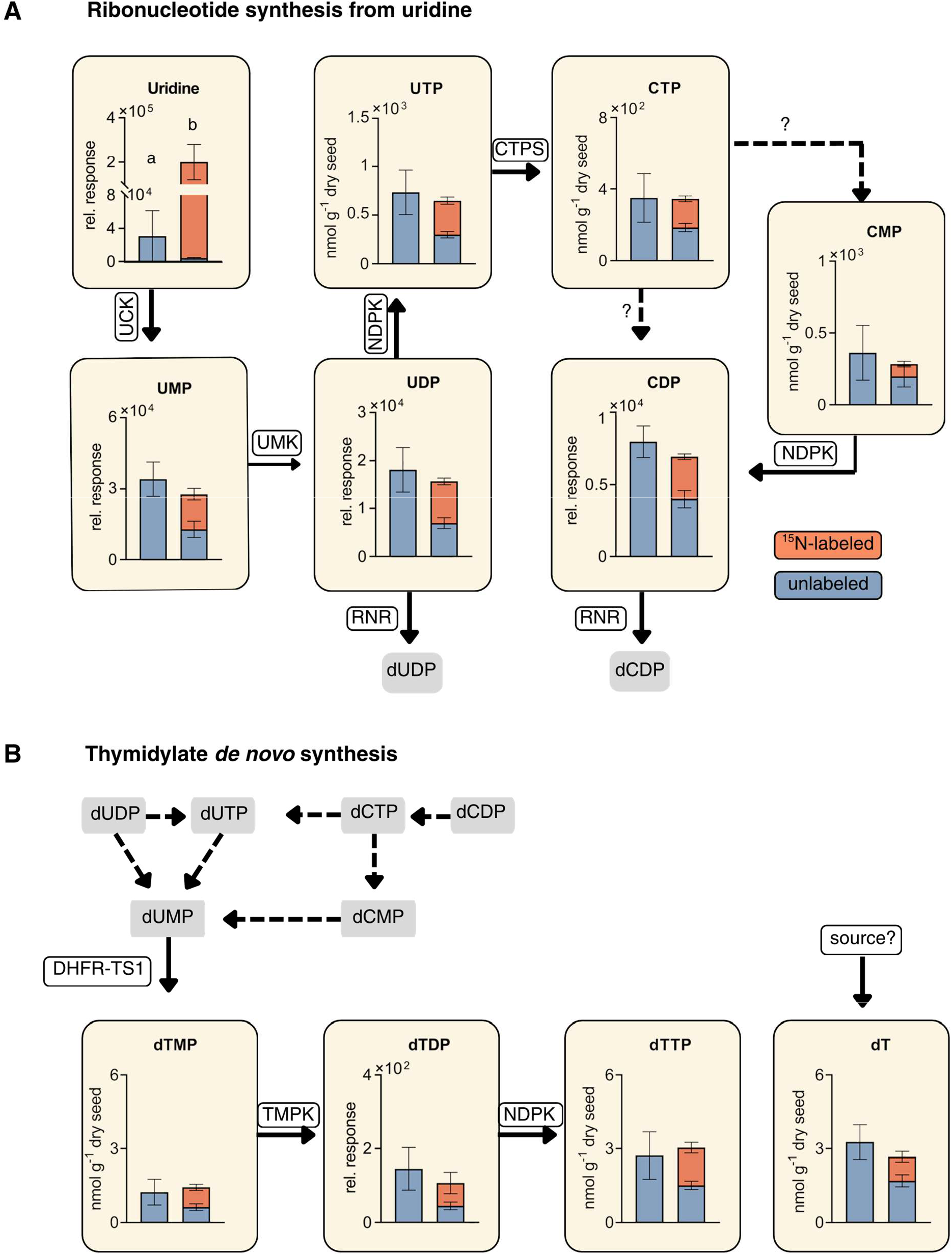
Pyrimidine ribonucleoside (rN), ribonucleotide (rNT) and thymidylate pools in germinating Arabidopsis seeds supplied with ¹⁵N-uridine. **(A)** Col-0 seeds were placed on ½ MS medium with or without ¹⁵N-uridine and harvested after 48 h. Stacked bars show concentrations or relative responses of the pyrimidine rN uridine and the rNTs UMP, UDP, CDP, UTP, CTP and CMP: unlabeled (blue, lower column part) and ¹⁵N-labeled metabolite species (orange, upper column part). For each metabolite, the left bar represents control conditions (no ¹⁵N-uridine) and the right bar treatment with ¹⁵N-uridine. Data are means ± SD (*n* = 4 - 5). Arrows indicate enzymatic steps in pyrimidine metabolism (UCK, UMK, CTPS, RNR, NDPK); dashed arrows and question marks indicate putative conversions. **(B)** dTMP, dTDP, dTTP and dT in the same samples as shown in A. “source?” indicates that there may be an unknown input into the dT pool. The scheme above shows the current model for the conversion of ribonucleotides into thymidylates. It is clear that finally dUMP is converted by DHFR–TS to dTMP followed by phosphorylation by TMPK and NDPK. However, how dUMP is generated is currently unclear. Dashed arrows indicate possible reactions.

In the labeled samples, the uridine pool was labeled to about 97% and was increased approximately tenfold compared to the unlabeled controls (Fig. 1A, Suppl. Table 1). However, this excess of uridine did not alter the total concentrations of other pyrimidines. In general, the label was distributed within the uridylate pool (UMP, UDP, and UTP; Fig. 1A) and UDP-Glucose (Suppl. Fig. 1) as well as within the cytidylate pool (CMP, CDP, and CTP; Fig. 1A). All these metabolite pools were labeled to approximately 50% except for CMP which was labeled only up to 31% on average. It seems that CTP produced by CTPS from UTP (Daumann et al. 2018; Witte and Herde 2020) can be dephosphorylated directly by a so far unknown CTP phosphatase giving rise to well labeled CDP whereas the lower label in CMP indicates that the alternative route to CDP via CMP is less prominent (Fig. 1A).

Having entered the dNT pool either as dUDP or dCDP, the label can only be incorporated into the thymidylate pool via the formation of dUMP and its subsequent methylation to dTMP (Gorelova et al. 2017) (Fig. 1B). We observed that approximately half of the dTMP (56%), dTDP (56%), and dTTP (50%) pools were labeled (Fig. 1 B, Suppl. Table 1). This shows that dNTPs are indeed synthesized during this period via *de novo* dNTP synthesis through RNR. The label was not only detected in the thymidylates, i.e. in phosphorylated dT, but also in dT itself (36%, Fig. 1B). This strongly suggests that *de novo* synthesis of thymidylates contributes to the increasing dT pool during germination.

### Blocking deoxypyrimidine synthesis confirms *de novo* thymidylates as a major source of dT during GSE

To further assess whether indeed labeled dT is derived from *de novo* thymidylate synthesis, we combined metabolic labeling with the application of inhibitors of *de novo* dNTP synthesis. First, 1 mM hydroxyurea (HU) was applied to inhibit RNR (Fig. 3) (Roa et al. 2009; Wang and Liu 2006). This resulted in reduced dATP, dCTP, and dGTP concentrations (Fig. 3A), as well as decreased labeled and unlabeled pools of dTTP, dTMP, and dT (Fig. 3B). RNR inhibition affected the unlabeled and labeled pools of dTMP (−68% and −63%) and dTTP (−68% and −55%) similarly, whereas the decrease in dT was greater in the labeled pool (−78%) than in the unlabeled pool (−36%). We next combined labeling with the application of 5-fluorouridine (5-FU), an inhibitor more specific to *de novo* thymidylate synthesis (Fig. 2, 4) (Kurasaka et al. 2022; Mori et al. 2022). 5-FU is converted to 5-fluoro-2’-deoxyuridine monophosphate (5F-dUMP) and inhibits the bifunctional DHFR-TS enzymes DHFR-TS1 in the cytosol and likely also DHFR-TS2 in mitochondria (Fig. 4A) (Corral et al. 2018; Gorelova et al. 2017; Kurasaka et al. 2022; Mori et al. 2022; Niehaus et al. 2022). Application of 50 µM 5-FU resulted in strongly reduced dTTP, dTMP, and dT concentrations. Only trace amounts of labeled dTMP and no labeled dT were detected in 5-FU-treated samples, supporting that label incorporation into dT depends on the *de novo* thymidylate synthesis pathway (Fig. 4A). The inhibition by 5-FU appears to have affected both DHFR-TS enzymes because it is known that in the absence of only one enzyme in the genetic mutants of *DHFR-TS1* or *DHFR-TS2* the respective thymidylate profiles are not markedly different from that of the wild type 48 h after transfer to growth conditions (Niehaus et al. 2022).

**Figure 2.**
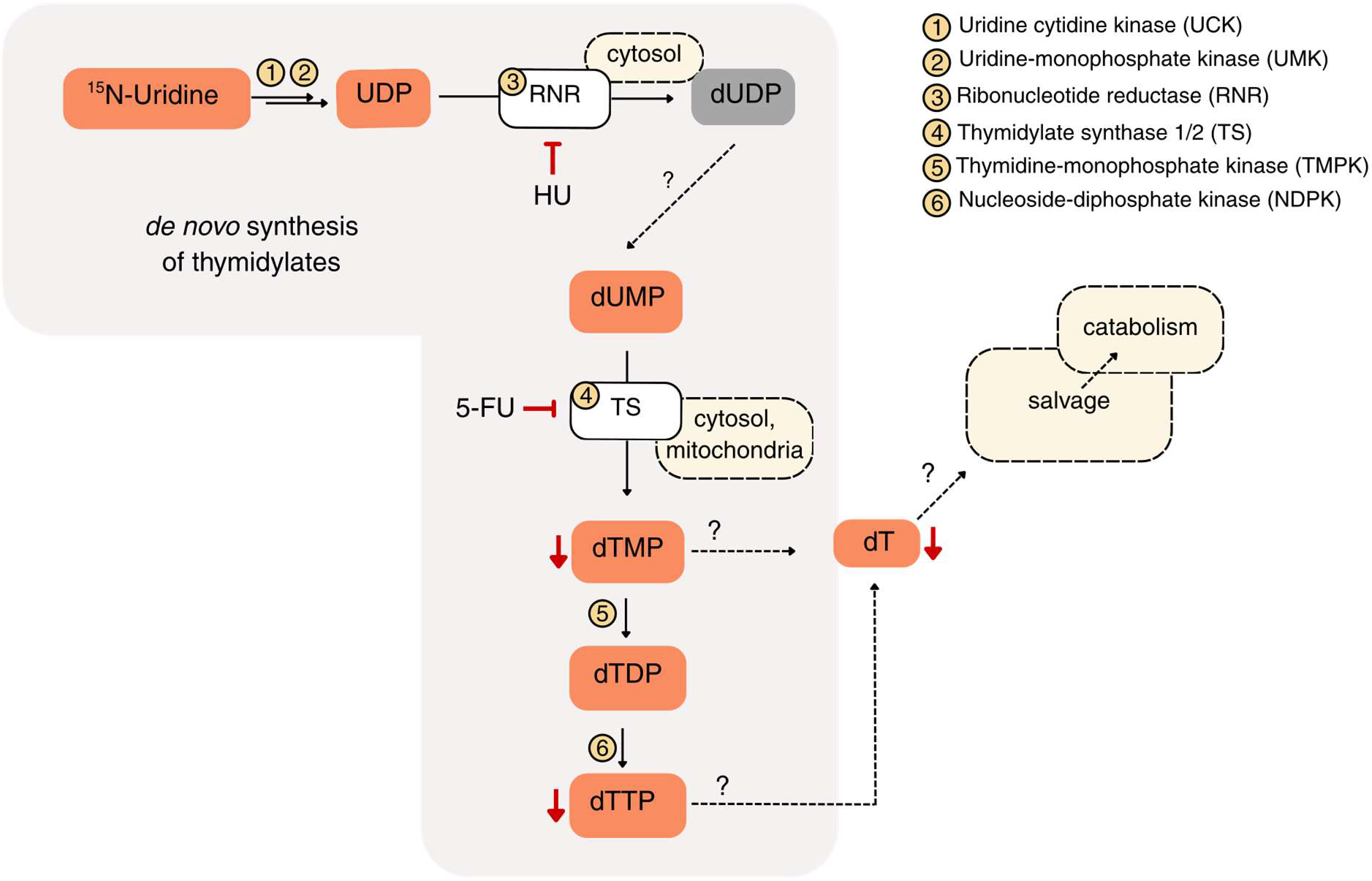
Schematic overview of ¹⁵N-uridine-derived *de novo* thymidylate synthesis and sites of inhibitor action. Pathway from ¹⁵N-uridine to thymidylate metabolites in germinating Arabidopsis seedlings. ¹⁵N-uridine is converted to UDP by uridine cytidine kinase (UCK, 1) and uridine-monophosphate kinase (UMK, 2). UDP is reduced to dUDP by ribonucleotide reductase (RNR, 3), which can be inhibited by hydroxyurea (HU). dUDP is further converted to dUMP and then to dTMP by thymidylate synthases 1 and 2 (TS, 4), which can be inhibited by 5-fluorouridine (5-FU). dTMP is phosphorylated to dTDP by thymidine-monophosphate kinase (TMPK, 5) and to dTTP by nucleoside-diphosphate kinase (NDPK, 6). Dashed arrows and question marks indicate steps that are not fully resolved, including the formation and further metabolism of deoxythymidine (dT). Red arrows highlight metabolites whose levels change in response to HU or 5-FU treatment.

**Figure 3.**
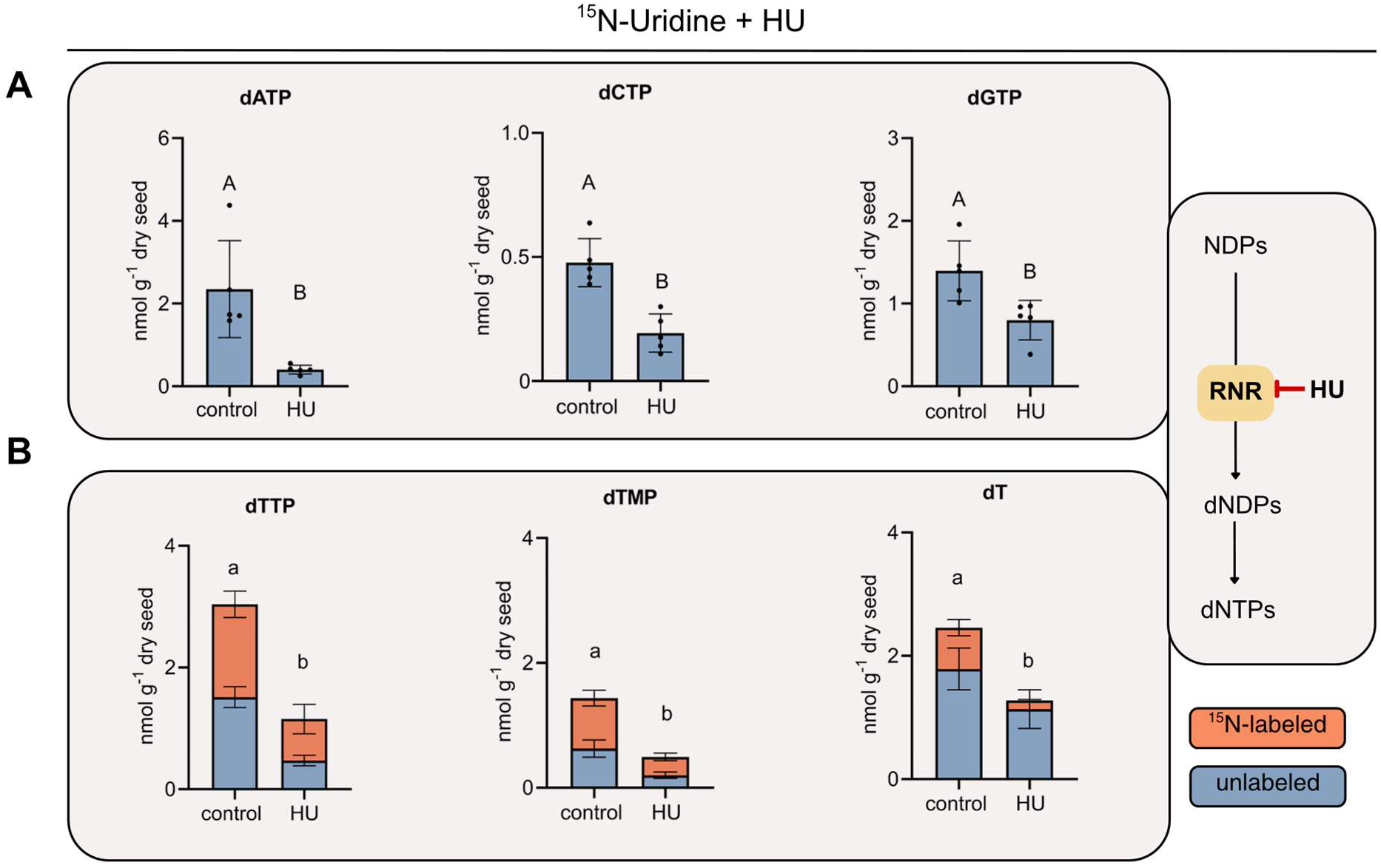
Effects of HU on dNT pools in germinating Arabidopsis seeds supplied with ¹⁵N-uridine. Col-0 seeds were grown on medium containing ¹⁵N-uridine and either none or 1 mM HU and were harvested 48 h after transfer to growth conditions. **(A)** dATP, dCTP and dGTP in seedlings treated with none or 1 mM HU. **(B)** dTTP, dTMP and dT in the same samples, shown as stacked bars: unlabeled (blue, lower column part) and ¹⁵N-labeled metabolite species (orange, upper column part). The scheme on the right illustrates inhibition of ribonucleotide reductase (RNR) by HU. Data are means ± SD (*n* = 5); different letters indicate statistically significant differences between treatments (P < 0.05); lower case letters indicate statistically significant differences in the unlabeled and labeled fraction.

**Figure 4.**
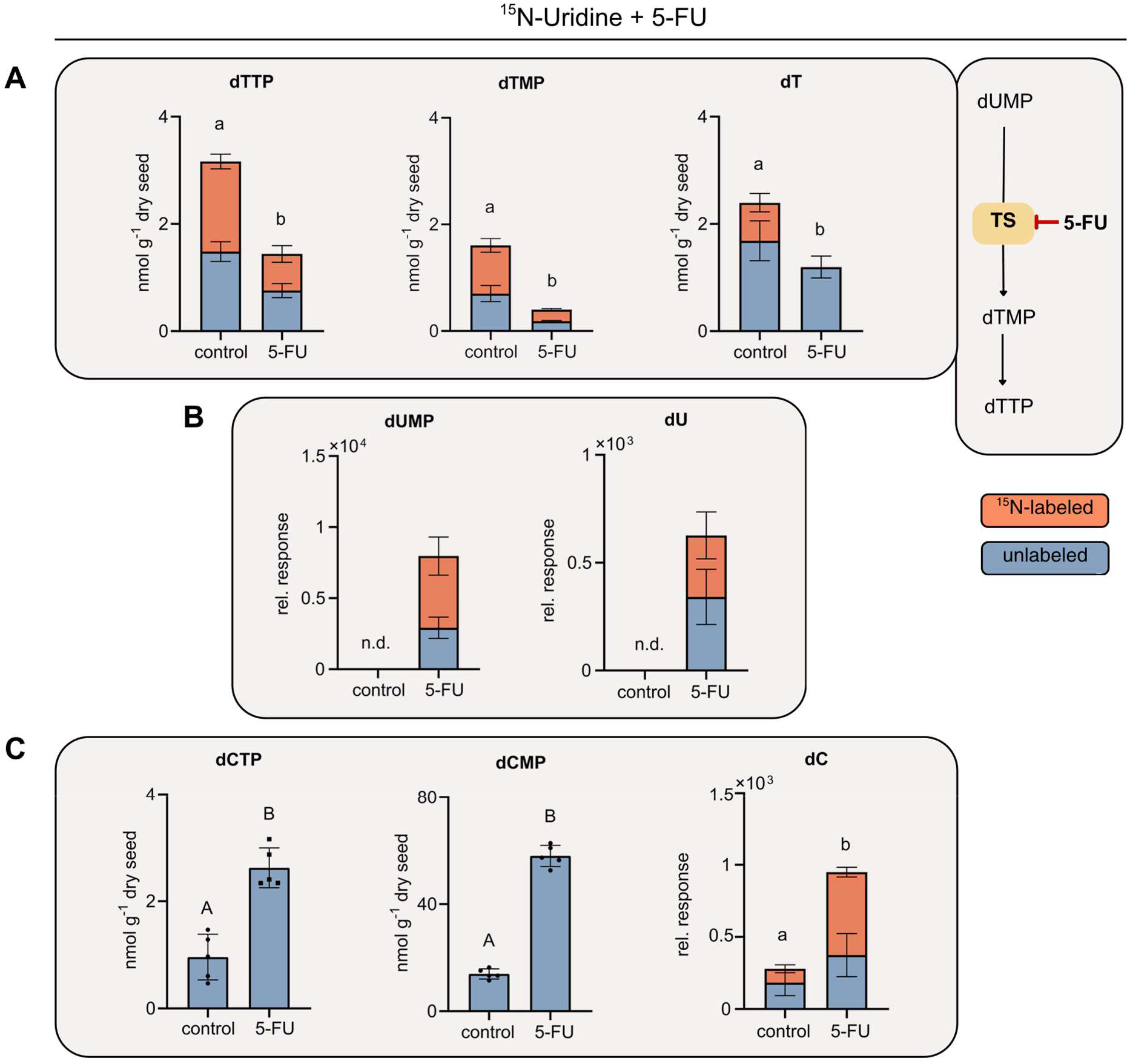
Effects of 5-FU on unlabeled and ¹⁵N-treated pyrimidines in germinating Arabidopsis seeds. Col-0 seeds were grown on medium containing ¹⁵N-uridine and either none or 50 µM 5-fluorouridine (5-FU) and harvested after 48 h after transfer to grwoth conditions. **(A)** dTTP, dTMP and dT in seedlings treated with none or 50 µM 5-FU, shown as stacked bars: unlabeled (blue, lower column part) and ¹⁵N-labeled metabolite species (orange, upper column part). **(B)** Relative LC–MS signals for unlabeled and ¹⁵N-labeled dUMP and dU in the same samples as shown in A; n.d. = not detectable **(C)** dCMP, dCTP and dC in the same samples as shown in A. The scheme on the right illustrates inhibition of thymidylate synthase (TS) by 5-FU. Data are means ± SD (*n* = 5); different letters indicate statistically significant differences between treatments (P < 0.05); lower case letters indicate statistically significant differences in the unlabeled and labeled fraction.

The DHFR-TS inhibition also led to an increase of dU and dUMP (Fig. 2, 4B) bringing them above the limit of detection (Suppl. Fig. 2A, B). Interestingly, both labeled and unlabeled dUMP and dU were detected, suggesting that the contribution of the *de novo* dTMP synthesis pathway to thymidylate and dT formation is greater than the proportion of incorporated label alone would indicate (Fig. 4B). This is expected, as ^15^N-uridine enters pyrimidine metabolism via salvage (Fig. 1A), while *de novo* ribonucleotide synthesis of UMP from orotate (Witte and Herde, 2020) likely continues to supply unlabeled UMP for deoxypyrimidine synthesis. Consistent with this notion not only the labeled but also the unlabeled dTTP and dTMP pools decrease upon HU and 5-FU treatment (Fig. 3B, Fig. 4A). Additionally, 5-FU-treated samples accumulated dCMP, dCTP, and dC (Fig. 4C). In dC, the labeled amount increased by 500% upon 5-FU (Fig. 4C). The analysis of dCMP and dCTP was restricted to the unlabeled metabolite species and samples because the labeled isotopologues could not be reliably quantified due to signal overlap with the internal isotope standard. The accumulation of dCMP (+316%) and dCTP (+173%) upon DHFR-TS inhibition indicates that the deamination of deoxycytidylates generally contributes to dUMP formation although pleiotropic effects of 5-FU on other enzymes than DHFR-TS could also be the cause of this accumulation (Marx and Alian 2015; Niu et al. 2017). In addition, dATP levels were significantly increased in 5-FU-treated samples, whereas dGTP remained unchanged (Suppl. Fig. 3).

Taken together, inhibitor treatments combined with ^15^N-uridine labeling confirm that *de novo* thymidylate synthesis is a major contributor to dT formation during GSE (Fig. 2).

### Loss of VEN4 drastically reduces dNs and depletes organellar DNA

The labeling and treatment with HU and 5-FU confirmed that *de novo* thymidylate synthesis contributes substantially to dT formation during GSE. This seems counterintuitive since *de novo* deoxypyrimidine synthesis produces phosphorylated thymidylates, which are first dephosphorylated to dT only to be then re-phosphorylated to dTTP in order to be used in DNA synthesis. So far, no plant enzyme has been identified that dephosphorylates dTTP to dT (Niehaus et al. 2022). A possible candidate is the *A. thaliana* enzyme VEN4 which is predicted to be a dNTP triphosphohydrolase. Loss of *VEN4* is associated with a chlorotic phenotype (Sarmiento-Mañús et al. 2023; Xu et al. 2020; Yoshida et al. 2018). To investigate the in vivo role of *VEN4*, we used a SALK T-DNA insertion line (SALK_031417). Plants homozygous for the T-DNA insertion were identified and designated *ven4-4* (Suppl. Fig. 4). A second line, *ven4-5*, was generated using CRISPR, yielding a plant with two homozygous single-base-pair insertions in *VEN4* that cause reading frame shifts, one in exon 2 and one in exon 5. From the segregating population, *ven4-5* plants lacking the transgene were selected and used for further characterization (Suppl. Fig. 5).

We observed the characteristic interveinal, chlorotic phenotype in both null mutants, particularly in the cotyledons and the first true leaves suggesting that there is a problem in chloroplast development (Fig. 5A) (Sarmiento-Mañús et al. 2023; Xu et al. 2020; Yoshida et al. 2018). The ratio of chloroplast and mitochondrial to nuclear DNA (cpDNA/ncDNA, mtDNA/ncDNA) was quantified in 48-h-old seedlings by qPCR as described in Niehaus et al. (2022). Both *VEN4* mutants showed significantly reduced cpDNA and mtDNA levels compared to the wild type. (Fig. 5B). Dry seeds of the *VEN4* mutants contained the same amount of organellar DNA as the wild type, demonstrating that the mutants do not have a deficit in cpDNA or mtDNA before germination (Suppl. Fig. 6) but that this deficit arises during the first 48 h of seedling development. The data show that efficient organellar DNA replication depends on VEN4.

**Figure 5.**
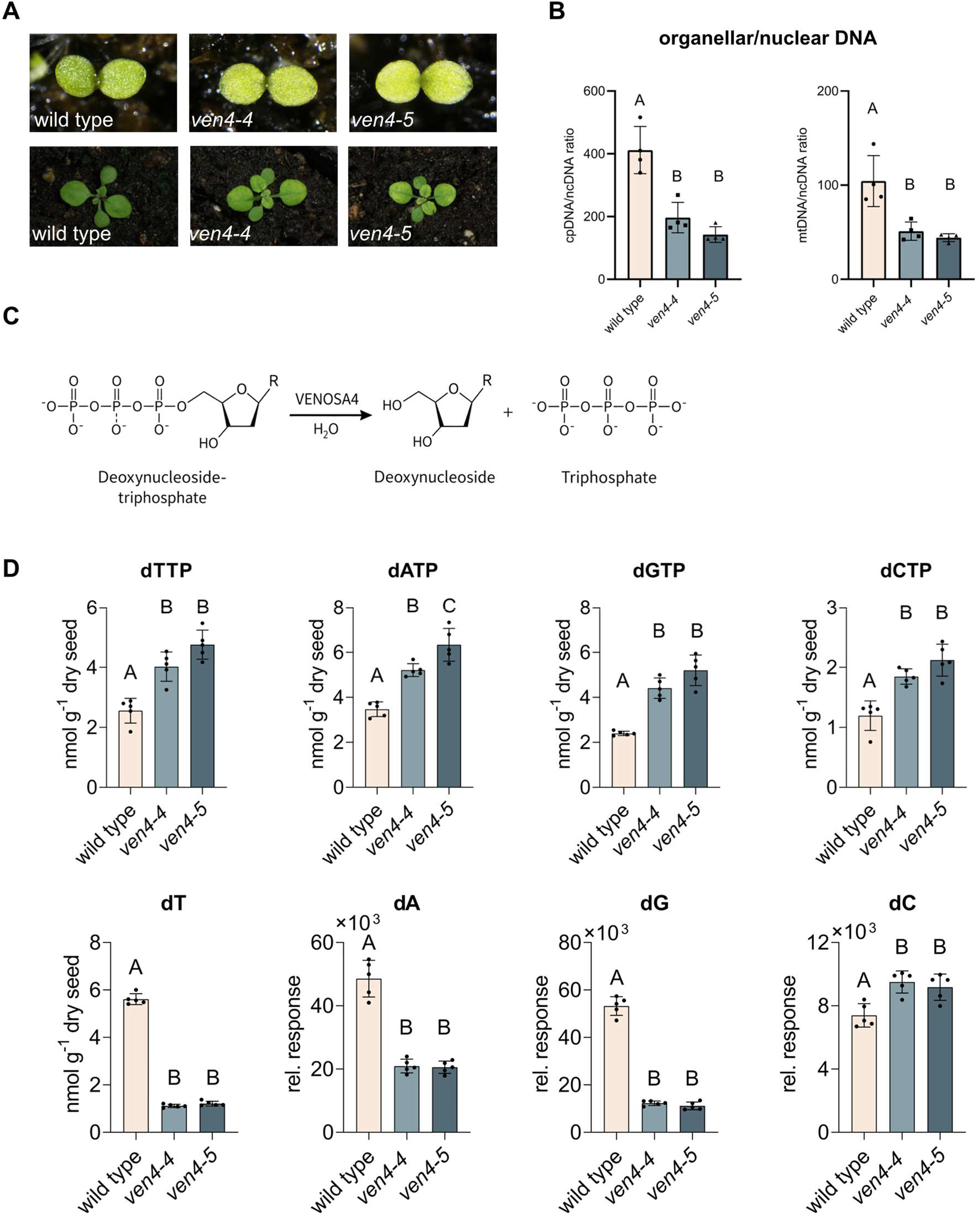
Organelle DNA abundance and dNT pools in *VEN4* mutants during GSE. **(A)** Representative images of wild type, *ven4-4* and *ven4-5* plants on soil 72 hours (top) and 21 days after transfer to growth conditions (bottom). **(B)** Organellar to nuclear DNA ratios (cpDNA/ncDNA and mtDNA/ncDNA) in wild type, *ven4-4* and *ven4-5*. Seeds were imbibed for 48 h at 4 °C in the dark, then moved to long-day conditions for 48 h before harvest. Organelle and nuclear DNA abundance were quantified by qPCR using primer pairs targeting the plastid gene RBCL (cpDNA), the mitochondrial gene COX1 (mtDNA) and the nuclear gene UBC21 (ncDNA). cpDNA/ncDNA and mtDNA/ncDNA ratios were derived from the corresponding Ct values. **(C)** Reaction scheme of VEN4-mediated dephosphorylation of dNTPs to the corresponding dNs. **(D)** dNTP pools (dTTP, dATP, dGTP and dCTP) and dN pools (dT, dA, dG and dC) in 48-hour-old wild type, *ven4-4* and *ven4-5*. Data are means ± SD (*n* = 4 - 5 biological replicates); different letters indicate statistically significant differences between genotypes (P < 0.05).

VEN4 shares a conserved histidine–aspartate (HD) phosphohydrolase domain with human SAMHD1 and has been shown to hydrolyze dGTP to dG *in vitro*. (Fig. 5C) (Lu et al. 2023). However, nucleotide concentrations have never been determined in *ven4* background to investigate the molecular *in vivo* effects of the mutation. We have quantified nucleotides by LC-MS/MS in the null mutants, *ven4-4* and *ven4-5*. Loss of VEN4 selectively affected dNTPs and dNs. All four canonical dNTPs accumulated in the mutant lines at about 50% to 120% above wild-type levels (Fig. 5D), whereas rNTP and rNMP amounts were unchanged (Suppl. Fig. 7A, B). The corresponding dNs were less abundant in the mutants, in particular dA, dG, and dT, but interestingly not dC (Fig. 5D). This accumulation of substrates and reduced concentration of products *in vivo* is consistent with proposed enzymatic function of VEN4 as dNTP triphosphohydrolase. The results strengthen our hypothesis that dT formation is strongly dependent on the *de novo* synthesis of thymidylates implicating VEN4 as main enzymatic player in this process.

### VEN4 couples *de novo*-derived dTTP to dT production

To assess whether VEN4 contributes to the conversion of thymidylates derived from *de novo* dNTP synthesis into dT we used the labeling approach which allows to metabolically distinguish *de novo* deoxypyrimidine synthesis-derived from salvage-derived thymidylates and dT. The two *VEN4* mutant lines were grown for 48 h on liquid ½ MS medium supplemented with ^15^N-uridine, and dNTP and dN levels were quantified. Both mutants again had significant less dT (by 75 and 71% in *ven4-4* and *ven4-5*, respectively) compared to the wild type, and accumulated dTTP (by 57% and 85%). The dTMP concentration was reduced by 71% and 68% compared to the wild type (Fig. 6A). Statistically significant changes (at P < 0.05) of total metabolite pool sizes are indicated by asterisks in Fig. 6. Furthermore, loss of *VEN4* affected the labeled dT pool more strongly than the unlabeled pool. Labeled dT was reduced by 87% and 85% in the mutant lines while unlabeled dT was reduced by 69% and 64%. Both pools were significantly (P < 0.05) smaller in the mutant, which is indicated by lower case letters in Fig. 6. Loss of *VEN4* also had a greater negative impact on the labeled (74% and 75%) than on the unlabeled (68% and 57 %) dTMP pool (Fig. 6). Interestingly, although there was more dTTP in the mutant lines, the labeled fraction of dTTP was not increased. This pattern is consistent with feedback downregulation of *de novo* dTTP synthesis by high dTTP levels, as described for the human deoxypyrimidine pathways (Hofer et al. 2012; Nordlund and Reichard 2006).

**Figure 6.**
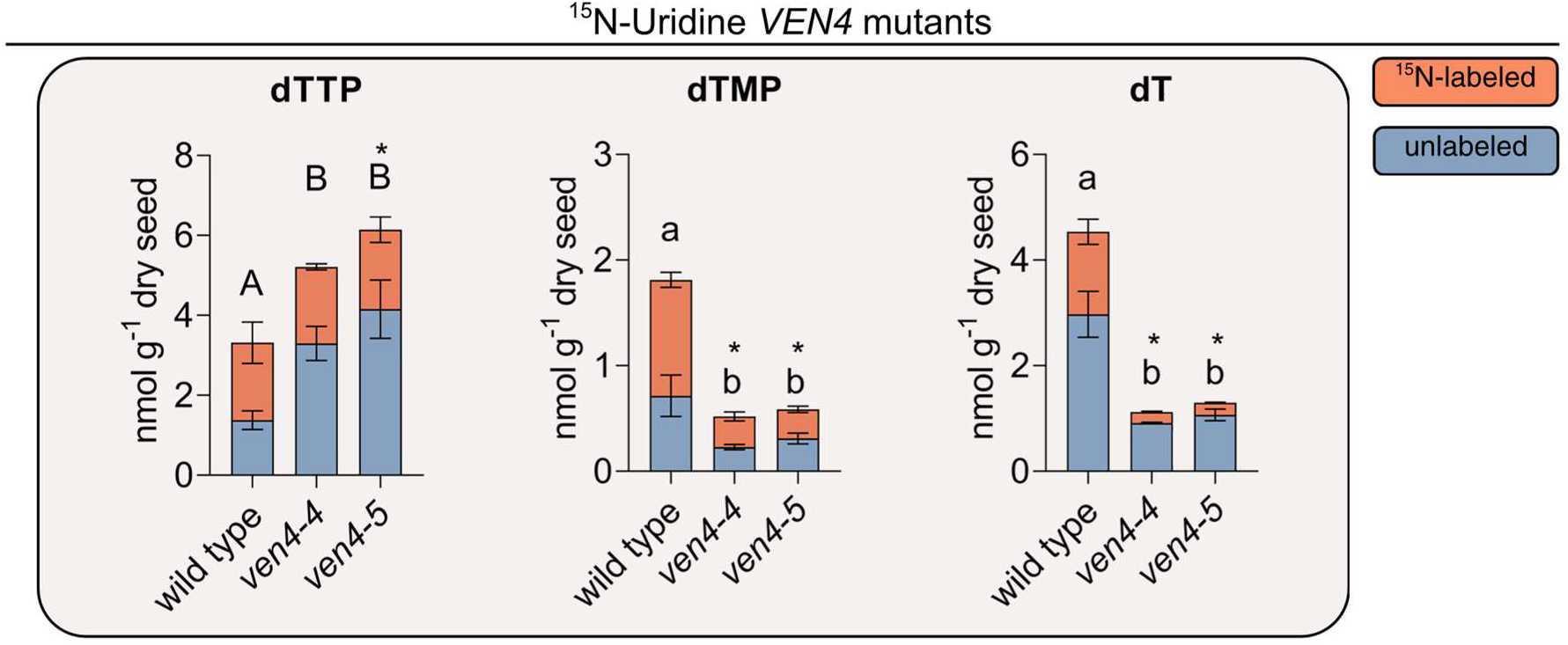
¹⁵N-uridine labeled thymidylates and dT in *VEN4* mutants. Wild type, *ven4-4* and *ven4-5* seeds were supplied with ¹⁵N-uridine. dTTP, dTMP and dT shown as stacked bars: unlabeled (blue) and ¹⁵N-labeled metabolite species (orange). Data are means ± SD (*n* = 3). Different letters indicate statistically significant differences (at P < 0.05) between genotypes in the unlabeled fraction; if letters are uppercase only the unlabeled fraction is different, if letters are lowercase unlabeled and labeled fractions are both different. Additionally, statistically significant differences in the total metabolite content (at P < 0.05) are indicated by asterisks.

Also, the lower concentration of dTMP (the first thymidylate product of *de novo* dTTP synthesis) in the mutants is consistent with such feedback regulation. However, the strong reduction of labeled dTMP in the mutants is striking and in stark contrast to the lack of effect on labeled dTTP in *ven4* background. This indicates that the labeled dTMP is mostly generated downstream of VEN4 (because it is influenced by *VEN4* mutation) consistent with the notion that a substantial fraction of dTMP is generated by re-phosphorylation of dT generated by VEN4, i.e., through the organellar salvage pathway via TK1b (Clausen et al. 2012; Le Ret et al. 2018; Niehaus et al. 2022). The data suggest that VEN4-derived dT is subsequently re-phosphorylated to dTMP in chloroplasts and mitochondria by TK1b and thereby fuels organellar DNA synthesis. Such a model could explain the counterintuitive observation that increased dNTP levels in the *VEN4* mutants coincide with reduced organellar DNA. It also links organellar dN salvage to cytosolic *de novo* dNTP synthesis and suggest a dependence of the former on the latter.

### *VEN4 - TK1b* genetic interaction and dN rescue link VEN4-dependent dN production to organellar salvage

The hypothesis, that the degradation of *de novo* synthesized dTTP to dT by VEN4 and the salvage of dT in the organelles might collectively contribute to cpDNA synthesis prompted us to investigate the relation of VEN4 and the corresponding dN salvage enzyme in *A. thaliana*, TK1b. TK1b, has been shown to localize to chloroplasts and mitochondria and to phosphorylate dT (Clausen et al. 2012; Niehaus et al. 2022; Pedroza-García et al. 2019). During GSE, loss of TK1b reduces the dTMP pool by about two thirds and leads to a smaller cpDNA/ncDNA ratio *in vivo* (Niehaus et al. 2022). Nevertheless, the *TK1b* mutant remains viable, because partially redundant pathways contribute to thymidylate synthesis. This provided an opportunity to further investigate the potential coupling between cytosolic dNTP degradation and organellar dN salvage. We generated a *VEN4 TK1b* double mutant (*ven4-4 tk1b*) by crossing. Heterozygous parental plants produced segregating progeny that included viable homozygous *VEN4 TK1b* double mutants. These homozygous plants completed their vegetative and reproductive development but did not produce viable seeds. The double mutant line had to be maintained through heterozygous plants, and homozygous double mutants used in the experiments were identified among their segregating progeny.

Seeds of Col-0, the two *VEN4* knockout lines, the *TK1b* knockout line, and the segregating *VEN4 TK1b* double knockout line were sown on soil and analyzed for cotyledon pigmentation. Cotyledons were assessed at 72, 96, and 144 h after transfer to long-day growth conditions (Fig. 7A). These time points were selected because the *tk1b* cotyledons are chlorotic directly after seedling establishment but become phenotypically indistinguishable from the wild type after five to seven days (Niehaus et al. 2022). The double mutant displayed a stronger phenotype than *tk1b* or *ven4* and was almost albino, showing a synergistic effect of *VEN4* and *TK1b* on cotyledon pigmentation (Fig. 7A). To objectively quantify the phenotype, the proportions of green, yellow, and white pixels in cotyledons were measured using Fiji (ImageJ) (Niehaus et al. 2022; Schindelin et al. 2012). At 72 hours, chlorosis and albinism were most pronounced in the mutants, with *tk1b* and *ven4-4 tk1b* showing the strongest phenotype (Fig. 7B). *Ven4-4* and *ven4-5* also exhibited a lower proportion of green pixels and a higher proportion of yellow pixels compared to Col-0. At 96 hours, *tk1b* still displayed mildly chlorotic cotyledons, whereas cotyledons of *ven4-4 tk1b* seedlings remained largely albinotic. The greening was restricted to meristematic and vascular (veinal) tissue in the double mutant, suggesting that the underlying metabolic processes may differ between these tissues and the mesophyll (Fig. 7A, B). After 144 h, all mutants except the double mutant were indistinguishable from Col-0 with respect to cotyledon pigmentation (Fig. 7B). The persistent phenotype of the double mutant might seem surprising as the single mutants recover. A possible explanation is the proposed mechanism underlying the phenotypic recovery of *tk1b*. It has been hypothesized that cytosolic TK1a partially compensates for the loss of organellar TK1b by phosphorylating dT to dTMP in the cytosol during early development implying some but limited thymidylate uptake from the cytosol by the organelles. This compensation becomes sufficient to restore the wild-type phenotype by 144 h in the cotyledons (Clausen et al. 2012; Niehaus et al. 2022; Pedroza-García et al. 2015; Pedroza-García et al. 2019). In *ven4* background, the cytosolic dT pool available to TK1a is strongly depleted (Fig. 5D), thereby limiting TK1a-mediated dTMP production and preventing phenotypic recovery of the double mutant.

**Figure 7.**
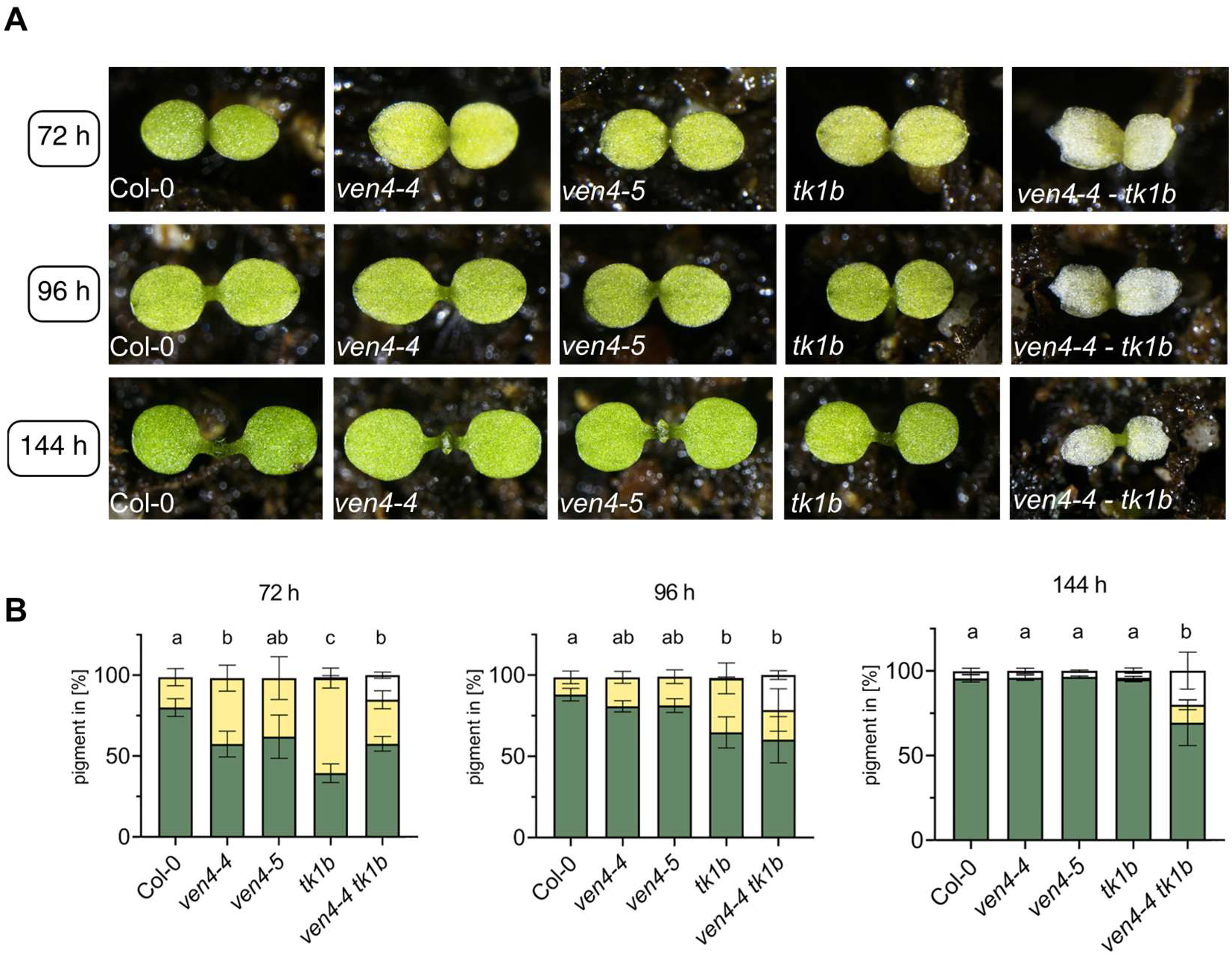
Phenotypic analysis of *tk1b, ven4* and *ven4 tk1b* seedlings during GSE. **(A)** Representative cotyledon images of Col-0, *ven4-4*, *ven4-5, tk1b* and *ven4-4 tk1b* grown on soil at 72 h, 96 h and 144 h after transfer to growth conditions. **(B)** Cotyledon pigment composition (green, yellow and white) at 72 h, 96 h and 144 h shown as stacked percentages. Data are means ± SD (*n* = 4 to 5 individual plants); different letters indicate statistically significant differences in percentage of green pigmentation in the cotyledons between genotypes (P < 0.05)

If the chlorotic phenotype of the *VEN4* mutants and the almost albinotic phenotype of the *VEN4 TK1b* double mutant result from insufficient dN supply to chloroplasts and mitochondria, exogenous dNs should alleviate these defects. To test this, we grew Col-0, the two *VEN4* knockout lines, *tk1b*, and *ven4-4 tk1b* for 14 days on solid ½ MS medium with or without either 100 µM dT or a mixture containing 100 µM each of dA, dG, dC, and dT (Fig. 8A). PlantCV-based image analysis was used to quantify total shoot area and the relative proportions of green, yellow, and white shoot tissue. Under control conditions, all mutants displayed reduced shoot areas and lower proportions of green shoot tissue than Col-0, with the *VEN4-4 TK1b* double mutant showing the most severe phenotype (Fig. 8B, C). The supplementation with dT alleviated the pigmentation defects of the single mutants reaching almost wild type levels (Fig. 8A, B). However, both *VEN4* mutants still had significantly smaller shoot areas than Col-0, showing that dT supplementation did not fully restore growth (Fig. 8C). In *tk1b*, dT restored shoot area to that of Col-0, although shoot pigmentation was only partially restored. Supplementation with all four dNs resulted in a more complete rescue of the single mutants. Here, *tk1b* and both *ven4* lines no longer differed from treatment-matched Col-0 in shoot area or in the proportions of green and yellow shoot tissue (Fig. 8B, C). The *VEN4-4 TK1b* double mutant showed a weaker response to dT and remained severely growth-retarded and largely albinotic. Treatment with all four dNs produced a stronger, although still incomplete, phenotypic improvement (Fig. 8B, C). The mean proportion of green shoot tissue increased from 0.6% under control conditions and 4.1% upon dT supplementation to 19.0% in the presence of all four dNs (Fig. 8B). Mean shoot area also increased from 3.2 mm² under control conditions and 5.6 mm² with dT to 11.5 mm² combined dN supplementation (Fig. 8C). Nevertheless, the double mutant remained significantly smaller and less green than the other genotypes.

**Figure 8.**
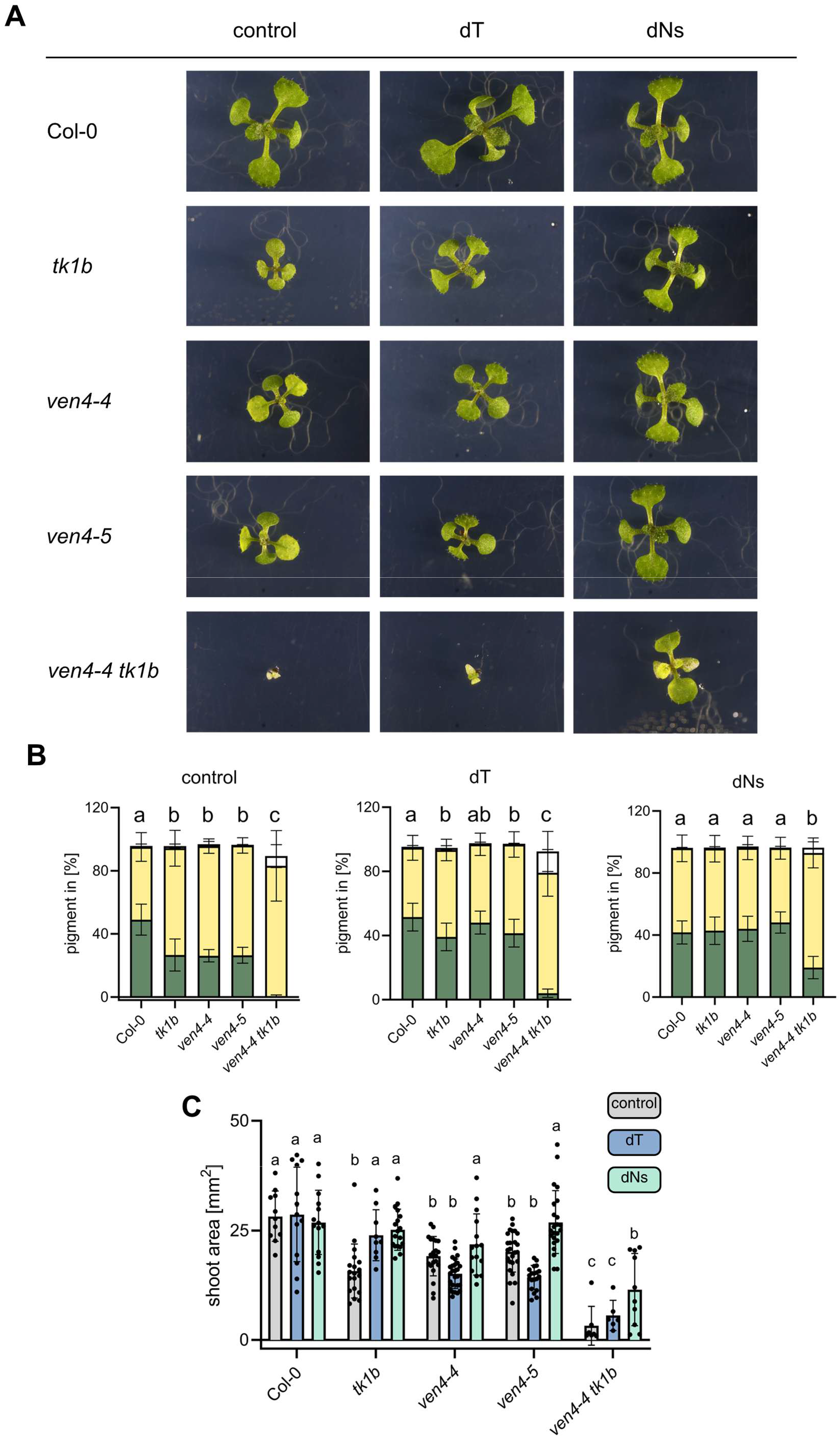
Phenotypic analysis of *ven4*, *tk1b* and *ven4-4 tk1b* under dN supplementation. Col-0, *tk1b*, *ven4-4*, *ven4-5* and *ven4-4 tk1b* seedlings were grown for 14 days on ½ MS agar medium without supplementation (control), with 100 µM dT, or with 100 µM each of the four canonical dNs (dA, dG, dC and dT). **(A)** Representative images of seedlings grown under the indicated supplementation conditions. **(B)** Shoot pigment composition shown as the relative proportions of green, yellow and white shoot tissue, quantified using PlantCV. **(C)** Total shoot area of seedlings grown under the indicated supplementation conditions, quantified using PlantCV. Data are means ± SD (*n* = 6 to 26 individual plants, see Suppl. Data set 1). Different letters indicate statistically significant differences between genotypes within the respective supplementation condition (P < 0.05). In (B), letters refer to the proportion of green shoot tissue.

Unlike the double mutant, *tk1b* retains VEN4-dependent production of dT, dA, and dG. The stronger response to all four dNs therefore suggests that VEN4-derived dNs other than dT also contribute to seedling development. These additional dNs likely enter salvage through deoxynucleoside kinase (dNK), while feeding dC may also contribute to thymidylate synthesis via dCMP and dUMP as outlined in the introduction (Niehaus et al. 2022; Niu et al. 2017).

This broader requirement for dNs may be particularly relevant to mitochondrial DNA precursor supply. Mitochondria can generate dTMP through DHFR-TS2 in the organelle but probably depend more strongly on salvage via dNK for the supply with dAMP, dGMP, and dCMP (Clausen et al. 2012; Clausen et al. 2014). The ^15^N-uridine feeding response (Fig. 6) together with the reduced mtDNA/ncDNA ratio in the *VEN4* mutants (Fig. 5B), suggests that an insufficient supply with dNs contributes to impaired mitochondrial DNA synthesis and reduced shoot growth in *ven4*. The genetic interaction between *VEN4-4* and *TK1b* and the differential response to dT and combined dN supplementation support a functional link between VEN4-dependent dN production and organellar dN phosphorylation.

### Depletion of labeled dTMP in *tk1b* supports TK1b-dependent phosphorylation of *de novo*-derived thymidine

To assess whether TK1b phosphorylates dT derived from *de novo*-synthesized thymidylates, we applied ¹⁵N-uridine labeling to the *TK1* mutants described by Niehaus et al. (2022). Both mutants showed slightly increased labeled and unlabeled dT concentrations. In *tk1a*, total and labeled dTMP were increased, while dTTP concentrations remained unchanged. The increase in dTMP despite loss of cytosolic TK1a suggests compensation by other routes of dTMP formation. One possibility is increased phosphorylation of the increased dT pool by TK1b, consistent with the proposed capacity of organellar dT salvage to compensate for impaired cytosolic salvage during GSE (Niehaus et al. 2022). Loss of *TK1b* resulted in a strong decrease in dTMP, including the labeled pool, while labeled dT was slightly elevated and total and labeled dTTP were increased compared with Col-0 (Fig. 9). Thus, dT remains available in *tk1b* but is less efficiently converted to dTMP. Our labeling experiments with *ven4* showed that a substantial fraction of labeled dT originates from VEN4-dependent dephosphorylation of *de novo*-derived dTTP. The accumulation of labeled dT together with the depletion of labeled dTMP in *tk1b* suggests that TK1b is operating downstream of VEN4-dependent dT formation. Genetic support for this functional link is provided by the severe phenotype of the *ven4-4 tk1b* double mutant (Fig. 7).

**Figure 9.**
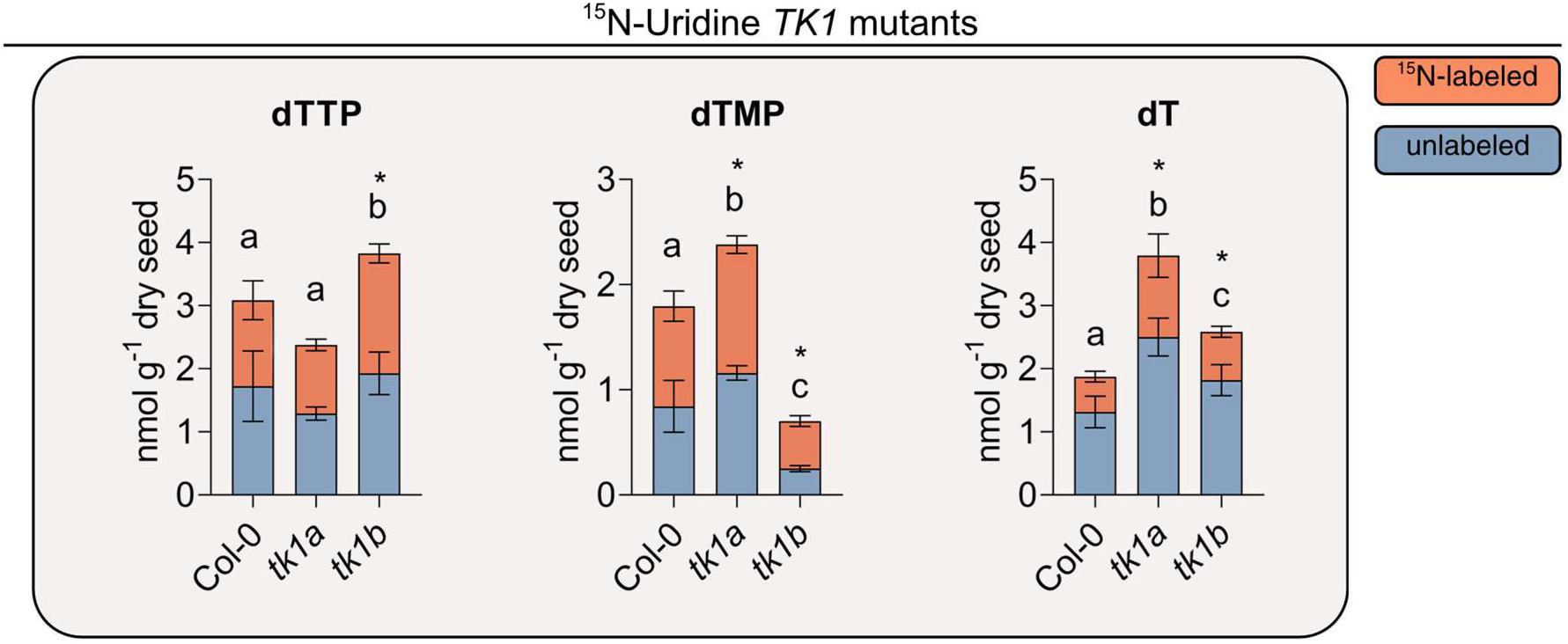
¹⁵N-uridine labeled thymidylates and dT in *TK1* mutants. Col-0, *tk1a* and *tk1b* seeds were supplied with ¹⁵N-uridine and harvested 48 h after transfer to growth conditions. Stacked bars show dTTP, dTMP and dT: unlabeled (blue) and ¹⁵N-labeled metabolite species (orange). Data are means ± SD (*n* = 5); different letters indicate statistically significant differences between genotypes in the unlabeled fraction; lower case letters indicate statistically significant differences in the unlabeled and labeled fraction; asterisks indicate statistically significant differences in total metabolite content (P < 0.05).

### Distinct cytosolic and mitochondrial contributions shape thymidylate supply and extend to later developmental stages

Although cpDNA and mtDNA were less abundant in *ven4* mutants, these DNAs could still be synthesized, indicating that alternative routes continue to supply precursors for organellar DNA synthesis. For thymidylates, two potentially compensatory routes are particularly plausible. First, the cytosolic TK1a could phosphorylate the residual dT to dTMP in the cytosol. This dTMP could contribute to chloroplast thymidylate supply, although the transport capacity for thymidylates across the chloroplast envelope appears to be limited during early GSE (Clausen et al. 2012; Niehaus et al. 2022; Pedroza-García et al. 2015; Pedroza-García et al. 2019). The accumulation of dT in *tk1a* is consistent with a contribution of TK1a to dT phosphorylation during GSE (Fig. 9) (Niehaus et al. 2022). Second, mitochondrial *de novo* dTMP synthesis through DHFR-TS2 could provide an additional source of thymidylates. Consistent with such a compensatory function, a *tk1b dhfr-ts2* double mutant generated by Niehaus et al. (2022) displayed more severe cotyledon chlorosis than *tk1b* and failed to recover fully, although the *DHFR-TS2* single mutant was phenotypically inconspicuous (Niehaus et al. 2022). The potential contributions of these pathways may also extend beyond early seedling establishment. Although the cotyledon phenotypes of *tk1b* and *ven4* became wild-type-like after six days, the *ven4* seedlings retained an interveinal chlorotic phenotype in true leaves even at 21 days after transfer to growth conditions (Fig. 10A). This indicates that VEN4-dependent dT/dN production remains important for chloroplast development even after the GSE phase. This is supported by the observation of disorganized chloroplast membrane structures in true leaves of 16-day-old Arabidopsis *VEN4* mutant plants (Sarmiento-Mañús et al. 2023).

**Figure 10.**
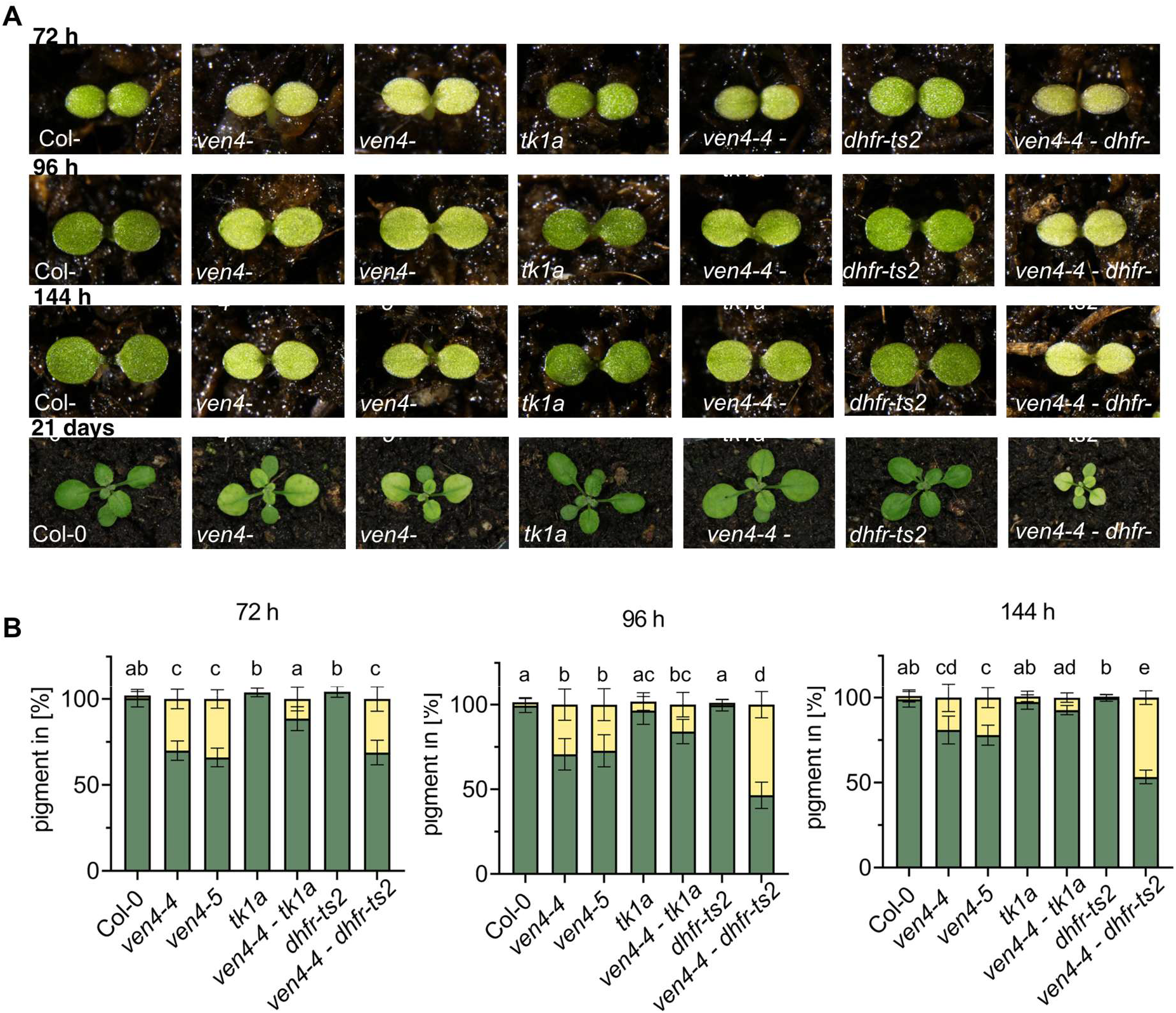
Phenotypic analysis of VEN4 and thymidine salvage or de novo thymidylate synthesis mutants. (A) Representative cotyledon images of Col-0, *ven4-4*, *ven4-5*, *tk1a*, *ven4-4 tk1a*, *dhfr-ts2* and *ven4-4 dhfr-ts2* grown on soil at 72 h, 96 h, 144 h and 21 days after transfer to growth conditions. (B) Cotyledon pigment composition (green and yellow) at 72 h, 96 h and 144 h shown as stacked percentages referenced to the mean of Col-0. Data are means ± SD (*n* = 3–5 individual plants); different letters indicate statistically significant differences in percentage of green pigmentation in the cotyledons between genotypes (P < 0.05).

To assess the contributions of cytosolic dT phosphorylation and mitochondrial dTMP synthesis in the *ven4* background, we generated *VEN4-4 TK1A* and *VEN4-4 DHFR-TS2* double mutants (Suppl. Fig. 8A, B, 9A, B). Individual seedlings were photographed, and cotyledon pigmentation was quantified using Fiji (ImageJ) at 72, 96, and 144 h after transfer to long-day growth conditions (Fig. 10A, B). As described above, the *VEN4* mutants displayed chlorotic cotyledons and interveinal chlorosis in true leaves, whereas the *TK1a* and *DHFR-TS2* single mutants could not be distinguished from Col-0 (Niehaus et al. 2022).

The *VEN4-4 TK1a* double mutant remained chlorotic relative to Col-0 but displayed a weaker phenotype than the *VEN4* single mutants, with chlorosis decreasing over the analyzed time course. Thus, loss of TK1a partially alleviated the *ven4*-associated phenotype. One possible explanation is that impaired cytosolic dT phosphorylation increases the availability of the remaining dT for uptake into chloroplasts and subsequent phosphorylation by TK1b. In the *ven4* background, TK1a activity therefore appears to be anti-compensatory, competing with TK1b for the limited dT pool (Figs. 9, 10).

Simultaneous disruption of mitochondrial *de novo* dTMP synthesis and VEN4-dependent dT generation in *dhfr-ts2 ven4-4* initially produced a chlorotic phenotype comparable to that of the *VEN4* single mutants (Fig. 10A, B). While the cotyledon phenotype of the *VEN4* mutants decreased over time, the cotyledons and true leaves of *dhfr-ts2 ven4-4* remained chlorotic. Interveinal chlorosis in adult *dhfr-ts2 ven4-4* plants also appeared more severe than in the *VEN4* single mutants. The persistent phenotype shows a mitochondrial contribution to thymidylate supply when VEN4-dependent dT production is impaired.

The opposite genetic interactions of *ven4-4* with *tk1a* and *dhfr-ts2* point to distinguished roles of these two pathways. TK1a activity seems to divert part of the remaining dT into the cytosolic thymidylate pool and thereby limit its availability for organellar salvage. DHFR-TS2 provides an alternative source of thymidylates that partially compensates for impaired VEN4-dependent dT production.

## Discussion

Since DNA replication in plants occurs in three genome-containing compartments, the question arises how plant cells ensure the provision of adequate amounts of DNA building blocks for the respective subcellular pools. Metabolically, dNTPs can be provided via two pathways, via the *de novo* synthesis of dNTPs in the cytosol and via the salvage of dNs in the cytosol and in the organelles (Clausen et al. 2012; Pedroza-García et al. 2019). Previous studies have shown that in particular dT salvage is of great importance for the replication of cpDNA during germination (Niehaus et al. 2022; Pedroza-García et al. 2019). However, the origin of the dT has remained unclear.

In this study, we combined ^15^N-uridine feeding in seedlings with the application of inhibitors of *de novo* dNTP and *de novo* dTMP synthesis and could thereby distinguish by LC-MS/MS between thymidylates formed from deoxypyrimidine *de novo* synthesis and salvage-derived thymidylates. In this way, we were able to trace the origin of a substantial proportion of dT back to *de novo* synthesis (Figs. 1B, 3B, 4A). Since the re-phosphorylation of dT by TK1b during germination is crucial for cpDNA synthesis (Niehaus et al. 2022), this finding implies a direct functional connection between *de novo* dTTP synthesis and organellar dT salvage. *De novo* synthesis and salvage are classically regarded as largely separate routes for dNTP supply, but our data change this view demonstrating that they are actually linearly connected in the same biosynthetic route during GSE. Thus, surprisingly, a main input for dN salvage enzymes comes from *de novo* synthesized dNTPs and not only from recycling of dNs derived from DNA turnover or repair processes as previously believed (Fig. 11).

**Figure 11.**
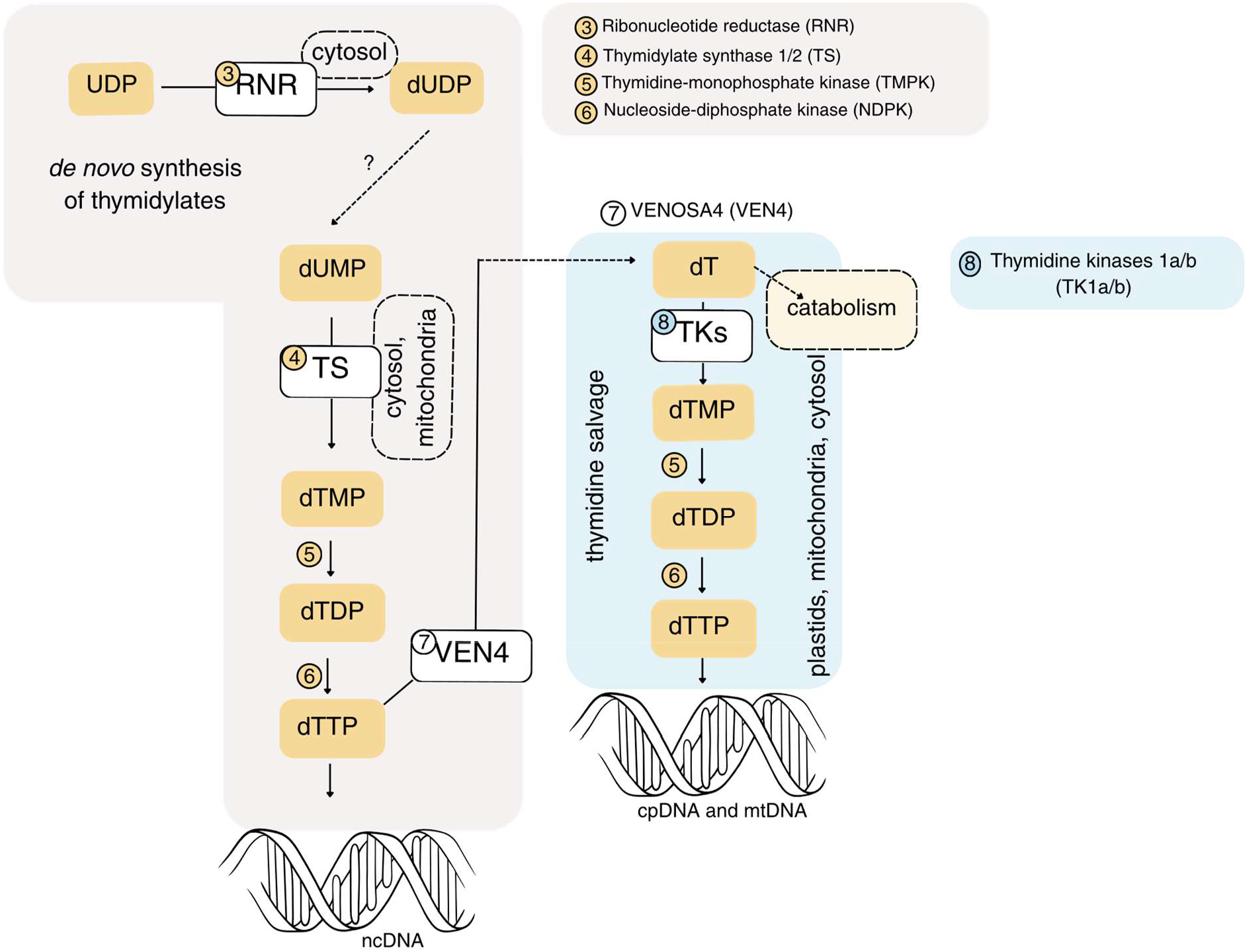
Integrative model of de novo thymidylate synthesis, VEN4 activity and dT salvage during GSE. Scheme of conversion of UDP into thymidylates via ribonucleotide reductase (RNR, 3), thymidylate synthase (TS, 4), thymidine-monophosphate kinase (TMPK, 5) and nucleoside-diphosphate kinase (NDPK, 6). VENOSA4 (VEN4, 7) dephosphorylates dTTP to dT, while thymidine kinases (TK1a/b; TKs, 8) convert dT back to dTMP as part of dT salvage in plastids, mitochondria and the cytosol. Dashed arrows and question marks indicate unresolved steps and potential links to catabolic routes.

We were able to identify VEN4 as the enzyme that connects *de novo* dNTP synthesis and dN salvage by its catabolic activity (Figs. 5, 8, 10). The analysis of the *VEN4* mutants strongly suggests that VEN4 is involved in the dephosphorylation of dNTPs and the resulting formation of dNs (Fig. 5D). For dTTP, uridine labeling shows clearly that VEN4 channels *de novo*-synthesized dTTP into the dT pool (Fig. 6). The nearly albino phenotype of *ven4-4 tk1b* provides genetic evidence for a functional interaction between the provision of dNs by VEN4 and their re-phosphorylation in the chloroplast (Fig. 7A, B). Consistent with this, exogenous addition of dNs can rescue the *ven4* phenotype (Fig. 8A-C). Together these findings demonstrate that VEN4 provides dNs from cytosolic *de novo* dNTP synthesis and that these become available for organellar salvage pathways. Interestingly, cpDNA and mtDNA synthesis depend greatly on the provision of dNs by VEN4 during GSE and probably also at later growth stages (Figs. 5B, 7A, B, 8A-C) indicating that the organelles are particularly competent for dN uptake and cannot be fully supported by cytosolic dNTs.

The metabolic function of VEN4 differs from that of its mammalian homolog SAMHD1 although both enzymes catalyze the same reaction. In animal cells, SAMHD1 is regarded as a dNTP-catabolizing enzyme that contributes substantially to the control and balancing of cellular dNTP pools (Franzolin et al. 2013; Goldstone et al. 2011). Conceptually, its function is to remove excess dNTPs. By contrast, VEN4 is part of a biosynthetic route for organellar dNTP supply. This explains the initially paradoxical observation that despite higher dNTP concentrations in *ven4*, the mutant has less cpDNA and mtDNA than the wild type (Fig. 5B, D). It becomes clear that for successful organellar DNA synthesis not only the total dNT concentrations are decisive, but in particular their phosphorylation state, localization, and availability to the respective compartment. In this respect the requirements of human cells are different since reduced SAMHD1 activity can rescue mitochondrial DNA deficiency in a mitochondrial dGK mutant (Franzolin et al. 2015). Thus, in the mammalian system, an increase in cytosolic dNTP availability can compensate for the loss of mitochondrial salvage (Franzolin et al. 2015), which is in stark contrast to what we have observed in Arabidopsis (Fig. 6B, D).

This different capacity of animal and plant cells to use cytosolic dNTs for organellar DNA synthesis points to differences in the organization of intracellular nucleotide transport between these organisms. Animal mitochondria have the ability to take up dNTs via members of the solute carrier 25 (SLC25) family which thereby contribute to mtDNA synthesis (Di Noia et al. 2014). Plants also possess numerous members of the mitochondrial SLC25/mitochondrial carrier family (MCF) but a function as dNT transporters has not been reported (Moller et al. 2020; Traba et al. 2011). Nonetheless, our data show that dNT transport into plant organelles exists but is not sufficient to fully support cpDNA and mtDNA synthesis.

Transport of dNs is also possible in mammalian cells via members of the Equilibrative Nucleoside Transporter (ENT) family, including mitochondrial ENT3 (Govindarajan et al. 2009). In quiescent cells, loss of mitochondrial salvage of dG or dT leads to mtDNA depletion in mammals (Franzolin et al. 2015; Villarroya et al. 2011) indicating that dN import is of significance in mammalian mitochondria. In the case of a dGK mutant, however, it was shown that this depletion can be compensated by increased cytosolic dNTP levels (Franzolin et al. 2015). Interestingly, depletion of ENT3 has been associated with an increased mtDNA amount in β-cells (Liu et al. 2015). The uptake of salvage substrates into mitochondria thus appears to be compensable by alternative pathways in mammals. Other plant species than Arabidopsis may be even more strongly dependent on organellar re-phosphorylation of dNs. In maize, a single dT kinase 1, CPTK1/ZmTK1, is dual-targeted to chloroplasts and mitochondria. Loss of CPTK1 causes albinism and a strong reduction in plastid genome copy number, demonstrating that local kinase capacity represents a substantial limitation for maintenance of the plastid genome in this species (Le Ret et al. 2018; Nájera-Martínez et al. 2020).

However, the partially stronger and longer-lasting chlorotic phenotype of the *ven4-4 dhfr-ts2* double mutant shows that a certain degree of metabolic independence of dN supply also exists in plant organelles, or at least in some species such as Arabidopsis (Fig. 10A, B). Mitochondrial DHFR-TS2 contributes only to a limited extent to the total dTMP pool during germination, which agrees with the wild type phenotype of the respective single mutant (Gorelova et al. 2017; Niehaus et al. 2022). In *ven4-4* background, however, the thymidylates provided by DHFR-TS2 clearly contribute to supporting the chloroplast, as loss of DHFR-TS2 results in a more persistent chlorotic phenotype in the *ven4-4* background (Fig. 10A, B).

Why do dNs play such an important role for the distribution of DNA precursors in plants? If organellar dN transport occured via an equilibrate, gradient-dependent mechanism, the direction of net flux would be determined by the concentration of the respective dN on both sides of the membrane (Boswell-Casteel and Hays 2017). Salvage of dNs in the chloroplast and mitochondria would generate dNTs and thereby trap them in the organelle (Jaehme and Slotboom 2015) and make them unavailable for nuclear DNA synthesis. Such trapping would work better for dNs because known transporters for (d)NTs can operate with different phosphorylation states and often exchange two (d)NT species over the membrane (Di Noia et al. 2014). In this case transport is not simply dependent on the transported target but also on the concentration of another nucleotide that would require regulation. In mitochondria of mouse livers, dTMP-specific transport has been demonstrated and speculated to rely on a uniport mechanism, but the identity of the transporter remains to be elucidated (Ferraro et al. 2006). In a recent review, MacVicar states that it remains unclear whether mammalian mitochondria have a preferred phosphorylation state in which they take up dN(T)s (MacVicar 2025). Equilibrative transport, in contrast, would allow for a direct feedback mechanism that couples demand and transport. When DNA synthesis rates decrease in the organelle, dNTP concentrations increase and salvage enzymes may be feedback inhibited. Such an inhibition by dNTPs has been described for TK1-family dN kinases, although it has so far not been directly demonstrated for Arabidopsis TK1b (Larsen et al. 2014; Welin et al. 2004). As consequence the free dN concentration in the organelle would rise and dN import cease or even be reversed to export. In the cytosol the dNs may then lead to product inhibition of VEN4 and in case of dT serve as substrate for TK1a, which makes them available for ncDNA synthesis. In this speculative model, VEN4-mediated dN production, organellar dN consumption by salvage and organellar DNA synthesis rate would be coordinated. Such a feedback architecture is an attractive concept but has not been experimentally tested so far.

## Material and Methods

### Plant material

*Arabidopsis thaliana* accession Columbia-0 (Col-0) served as the wild-type background. A segregating T-DNA insertion line from the SALK collection (*ven4-4* SALK_031417; Alonso et al., 2003) was acquired from the Nottingham Arabidopsis Stock Centre. A homozygous knockout line and a corresponding WT were selected from the segregating population. Arabidopsis T-DNA insertion mutants (*ven4-4*) and respective wild-type plants were identified by PCR using the following primer combinations: P1477 and P2572 (wild-type reaction); P2572 and P1316 (T-DNA reaction). The PCR product of the T-DNA reaction was sent for sequencing. A second knockout line was obtained via genome editing (*ven4-5*) according to (Rinne et al. 2021). The guide RNAs were selected according to CRISPR-P 2.0 (Liu et al., 2017). Editing events were screened using capillary sequence, as described recently Rinne et al., (2021), and confirmed by sequencing.

We generated *ven4-4 tk1a*, *ven4-4 tk1b*, and *ven4-4 dhfr-ts2* by crossing of the respective single mutants. Double mutants were identified in segregating progeny by PCR (Suppl. Data Set 1). Homozygous *ven4-4 tk1b* plants did not produce viable seeds. Therefore, we maintained this line through heterozygous plants and identified homozygous double mutants among segregating progeny. We grew all mutant lines and their Col-0 or segregating wild-type controls in parallel to obtain uniform seed batches. Seeds used for cotyledon pigmentation analysis in Fig. 7 were stored for 6 months after harvest, whereas seeds used in Fig. 10 were stored for 1 month after harvest, which may account for the stronger chlorotic cotyledon phenotype of *ven4-4* and *ven4-5* in Fig. 10.

### Growth conditions Soil-grown plants

Seeds were surface-sterilized with 70% (v/v) ethanol for 10 min, dried, and sown on soil (Einheitserde Special, Werkverband e.V.). After 48 h stratification at 4°C in darkness, plants were transferred to long-day conditions (16 h light/8 h dark; 100 µmol m⁻² s⁻¹; Osram Fluora 36W/77; 22°C day/20°C night; 60% relative humidity) and grown for the indicated duration.

### Seedlings on liquid ½ MS

Seeds were surface-sterilized with 70% (v/v) ethanol for 10 min, dried, and divided into 10 ± 0.5 mg aliquots. Two layers of autoclaved filter paper (6 × 6 cm) were placed in Petri dishes (94 × 16 mm) and soaked with 2 mL cold, sterile, modified liquid ½ MS medium(Niehaus et al. 2022). For isotope-labeling experiments, ¹⁵N₂-uridine (Eurisotop, NUM-812-25) was added to 250 µM. For inhibitor experiments, hydroxyurea (HU; Abcam, ab142613) or 5-fluorouridine (5-FU; Thermo Scientific, A52083.06) was added to 1 mM or 50 µM, respectively, either to unlabeled medium or to medium containing 250 µM ¹⁵N₂-uridine. Seeds were distributed evenly on the filter paper, stratified for 48 h at 4°C in darkness, and then transferred to the long-day conditions described above. Seedlings were harvested after 48 h(Niehaus et al. 2022). The modified ½ MS medium contained 3 mM CaCl₂, 1.5 mM MgSO₄, 1.25 mM KH₂PO₄, 18.7 mM KNO₃, 0.1 mM FeSO₄, 0.1 mM Na₂EDTA, 0.13 mM MnSO₄, 0.1 mM H₃BO₃, 0.03 mM ZnSO₄, 1 µM Na₂MoO₄, 0.1 µM CuSO₄, 0.1 µM NiCl₂, and 0.5 g L⁻¹ MES. The pH was adjusted to 5.7 with KOH.

### Seedlings on solid ½ MS

Seeds were grown for 14 days on the modified ½ MS medium described above, solidified with 0.8% (w/v) agar. The medium contained either no deoxyribonucleoside supplement (control), 100 µM dT, or 100 µM each of dA, dG, dC, and dT. Square Petri dishes (120 × 120 mm) were used and were kept in the dark at 4°C for 48 h before transfer to long-day conditions (16 h light/8 h dark; 100 µmol m⁻² s⁻¹; poly klima True Daylight PLUS White-LED; 22°C day/20°C night; 70% relative humidity). Plates were placed horizontally for growth.

### DNA isolation

Genomic DNA was isolated from 48-h-old seedlings and dry seeds. Each biological replicate contained 50 mg plant material. Seedlings were flash-frozen in liquid nitrogen before homogenization. Dry seeds were processed directly. Both sample types were homogenized in a bead mill at 16 Hz for 4 min and extracted with the same CTAB-based protocol, including RNase digestion, as described by (Niehaus et al. 2022)

### qPCR analysis

qPCR was performed according to (Niehaus et al. 2022)on a QuantStudio 3 Real-Time PCR System (Thermo Fisher Scientific) with qPCRBIO SyGreen Mix (PCR Biosystems) according to the manufacturer’s instructions. Reactions had a final volume of 10 µL and were run for 3 min at 95°C, followed by 40 cycles of 5 s at 95°C and 30 s at 60°C. Melt curves were recorded by heating to 95°C for 15 s, holding at 60°C for 1 min, and ramping to 95°C at 0.1°C s⁻¹. All products showed a single melting peak.

Each genotype comprised four to five biological replicates measured in three technical replicates. The same workflow and replicate structure were used for dry seeds. Ct values of organellar targets were normalized to the nuclear reference locus, and cpDNA/ncDNA and mtDNA/ncDNA ratios were calculated using the 2⁻ΔCt method (Livak and Schmittgen 2001). RBCL (AtCG00490), COX1 (AtMG01360), and UBC21 (At5G25760) were used as chloroplast, mitochondrial, and nuclear targets, respectively. Primer sequences are listed in Suppl. Data Set 1.

### Cotyledon image processing and pixel-based pigmentation quantification

Cotyledons of soil grown seedlings were imaged at 72, 96, and 144 h after transfer to long-day conditions using a Nikon SMZ25 stereo microscope equipped with a Nikon DS-Ri2 camera (Nikon, Minato, Japan) at 4,908 × 3,264 pixels. Microscope and illumination settings were kept constant across samples.

Pigmentation was quantified in Fiji/ImageJ with a two-step workflow according to (Niehaus et al. 2022)with minor modifications. A rectangular selection was drawn around the cotyledons, and a custom macro generated a centered 1,000 × 1,000-pixel crop. The crop was duplicated, contrast was enhanced (Enhance Contrast; saturated = 0.35%), a sharpening filter was applied, and the image was saved as TIFF. A second macro converted images to HSB color space and classified pixels as white/pale (Saturation ≤ 15 and Brightness ≥ 200), green (Hue 46–100), or yellow (Hue 35–45). Percentages were calculated from the sum of classified pixels. For Fig. 10, saturation was kept at 0–255 and brightness was set automatically; pixels were classified only as green (Hue 46–100) or yellow (Hue 35–45) and are reported as percentages of the mean of Col-0.

### PlantCV-based image processing and phenotypic quantification

Fourteen-day-old plate-grown seedlings were imaged with the same Nikon SMZ25/DS-Ri2 setup. Images were acquired in NIS-Elements BR with an SHR Plan Apo 0.5× objective, 1.5× zoom, a resolution of 4,908 × 3,264 pixels (3 × 8-bit RGB), continuous uniform illumination, a fixed exposure time of 500 ms, and an analog gain of 2.0×. The image calibration was 3.90 µm pixel⁻¹. As the plate setup included a more complex background due to visible roots and the more complex architecture of the rosettes, we used a custom PlantCV workflow in Python 3.12 (PlantCV version 4.11; plantcv_feeding_workflow.py) to analyze images of fourteen-day-old plants. HSV and LAB channels were used to generate a broad shoot mask that included green, chlorotic, and pale/bleached tissue. The shoot mask used H = 17–63, S ≥ 31, and V ≥ 46; pale tissue was additionally detected with S ≤ 76, V ≥ 107, and LAB-b ≥ 128. The combined mask was refined using standard morphological operations to remove small artifacts, close small gaps, and fill holes within segmented shoot regions. Components smaller than 80 pixels and root-like components with elongation > 8 and fill < 0.25 were removed. Within the shoot mask, pixels were classified as green (H = 34–63, S ≥ 41, V ≥ 46), yellow/chlorotic (H = 16–<34, S ≥ 26, V ≥ 51), white/bleached (S ≤ 38, V ≥ 135), or other. The same thresholds were used for all genotypes and treatments. Total shoot area and the relative proportions of green, yellow/chlorotic, and white/bleached tissue were calculated for each seedling. Areas were converted to mm² using the 3.90 µm pixel⁻¹ calibration.

All segmentations were checked using automatically generated QC overlays and group-wise montages. Images were excluded for severe condensation, extensive air bubbles or reflections, poor focus, truncated shoots, or unresolvable overlap.

### Metabolite analysis

Seedlings were collected from the filter paper into 2-mL Safe-Lock tubes containing five 2-mm steel beads and one 6.5-mm steel bead, flash-frozen in liquid nitrogen, and stored at −80°C or processed immediately. Frozen material was bead-milled twice for 4 min at 16 Hz with liquid-nitrogen cooling between runs. Sample preparation by solid-phase extraction (SPE), the LC-MS/MS setup, and isotope-dilution quantification followed (Straube et al. 2023). The chromatographic gradients were modified as described below. Both nucleotide and nucleoside analyses were performed on an Agilent 1290 Infinity II LC coupled to an Agilent 6470 triple quadrupole MS.

Nucleotides were separated on a Hypercarb column (50 × 4.6 mm, 5 µm; Thermo Scientific) with mobile phase A (5 mM ammonium acetate, pH 9.5, in water) and B (acetonitrile) using the following gradient: 0–3 min, 100% A; 18 min, 70% A; 19–22 min, 0% A; 22.5 min, return to 100% A; re-equilibration at 100% A until 30 min. Chromatography was run at 0.6 mL min⁻¹ and 35°C with a 10 µL injection.

Nucleosides were separated on a Polaris C18-A column (4.6 × 50 mm, 3 µm) at 0.6 mL min⁻¹ and 30°C with a 10 µL injection. Mobile phases were 0.0075% (v/v) formic acid in water (A) and 0.0075% (v/v) formic acid in methanol (B). The gradient was 96% A at 0 min, 35% A at 8.00 min, 100% A from 8.20 to 10.00 min, and 96% A from 10.10 to 12.50 min. Compounds were detected in positive MRM mode. Transitions and fragmentor settings for unlabeled analytes followed Straube et al. (2021); additional transitions with the corresponding mass shifts were used for isotopically labeled metabolites. For dCMP and dCTP, labeled isotopologues were not quantified because their signals overlapped with the internal standard. Metabolite concentrations were normalized to the dry weight of the starting material as described by (Niehaus et al. 2022).

### Statistical analysis

Data were analyzed in R 4.5.1 using RStudio 2025.05.1 and the CRAN packages multcomp and sandwich, following (Heinemann et al. 2021). We used two-sided Tukey-adjusted pairwise comparisons and a sandwich variance estimator to account for heteroscedasticity (Herberich et al. 2010; Hothorn et al. 2008). Adjusted P-values are provided in Suppl. Data Set 1.

## Supporting information

Supplementary Data Set 1

## Acknowledgements

We are grateful to Prof. Dr. Thomas Pfannschmidt for his valuable support and helpful discussions.

## Author contributions

MH and LF designed the study. LF performed the metabolomics experiments and mutant characterization, analyzed and interpreted the data, and wrote the first draft of the manuscript. CPW, MH, and HS edited the manuscript. HS designed and generated the CRISPR mutant. HT performed the genetic crosses and generated the double mutants used in this study. NP established the labeling procedure. All authors contributed to the final version of the manuscript and approved it for publication.

## Funding

This work was supported by Deutsche Forschungsgemeinschaft (DFG), HE5949/3-3.

## Competing interests

None declared.

## Data availability

The data that support the findings of this study are available in the Supporting Information of this article.

## Notes

### Competing Interest Statement

The authors have declared no competing interest.

## References

Bölter B, Sharma R, Soll J (2007). Localisation of Arabidopsis NDPK2—revisited. Planta 226(4): 1059–1065. DOI: 10.1007/s00425-007-0549-4.

Boswell-Casteel RC, Hays FA (2017). Equilibrative Nucleoside Transporters - A Review. Nucleosides, Nucleotides & Nucleic Acids 36(1): 7–30. DOI: 10.1080/15257770.2016.1210805.

Chen KL, Xu MX, Li GY, Liang H, Xia ZL, Liu X, et al. (2006). Identification of AtENT3 as the main transporter for uridine uptake in Arabidopsis roots. Cell Research 16(4): 377–388. DOI: 10.1038/sj.cr.7310049.

Clausen AR, Girandon L, Ali A, Knecht W, Rozpedowska E, Sandrini MPB, et al. (2012). Two thymidine kinases and one multisubstrate deoxyribonucleoside kinase salvage DNA precursors in Arabidopsis thaliana. The FEBS Journal 279(20): 3889–3897. DOI: 10.1111/j.1742-4658.2012.08747.x.

Clausen AR, Mutahir Z, Munch-Petersen B, Piškur J (2014). Plants Salvage Deoxyribonucleosides in Mitochondria. Nucleosides, Nucleotides & Nucleic Acids 33(4–6): 291–295. DOI: 10.1080/15257770.2013.853782.

Corral MG, Haywood J, Stehl LH, Stubbs KA, Murcha MW, Mylne JS (2018). Targeting plant DIHYDROFOLATE REDUCTASE with antifolates and mechanisms for genetic resistance. The Plant Journal 95(4): 727–742. DOI: 10.1111/tpj.13983.

Daumann M, Hickl D, Zimmer D, DeTar RA, Kunz HH, Möhlmann T (2018). Characterization of filament-forming CTP synthases from Arabidopsis thaliana. The Plant Journal 96(2): 316– 328. DOI: 10.1111/tpj.14032.

Di Noia MA, Todisco S, Cirigliano A, Rinaldi T, Agrimi G, Iacobazzi V, et al. (2014). The Human SLC25A33 and SLC25A36 Genes of Solute Carrier Family 25 Encode Two Mitochondrial Pyrimidine Nucleotide Transporters. Journal of Biological Chemistry 289(48): 33137– 33148. DOI: 10.1074/jbc.M114.610808.

Dorion S, Matton DP, Rivoal J (2006). Characterization of a cytosolic nucleoside diphosphate kinase associated with cell division and growth in potato. Planta 224(1): 108–124. DOI: 10.1007/s00425-005-0199-3.

Draper CK, Hays JB (2000). Replication of chloroplast, mitochondrial and nuclear DNA during growth of unirradiated and UVB-irradiated Arabidopsis leaves. The Plant Journal 23(2): 255–265. DOI: 10.1046/j.1365-313x.2000.00776.x.

Dubois E, Córdoba-Cañero D, Massot S, Siaud N, Gakière B, Domenichini S, et al. (2011). Homologous Recombination Is Stimulated by a Decrease in dUTPase in Arabidopsis. PLoS ONE 6(4): e18658. DOI: 10.1371/journal.pone.0018658.

Ferraro P, Nicolosi L, Bernardi P, Reichard P, Bianchi V (2006). Mitochondrial deoxynucleotide pool sizes in mouse liver and evidence for a transport mechanism for thymidine monophosphate. Proceedings of the National Academy of Sciences of the United States of America 103(49): 18586–18591. DOI: 10.1073/pnas.0609020103.

Franzolin E, Pontarin G, Rampazzo C, Miazzi C, Ferraro P, Palumbo E, et al. (2013). The deoxynucleotide triphosphohydrolase SAMHD1 is a major regulator of DNA precursor pools in mammalian cells. Proceedings of the National Academy of Sciences of the United States of America 110(35): 14272–14277. DOI: 10.1073/pnas.1312033110.

Franzolin E, Salata C, Bianchi V, Rampazzo C (2015). The Deoxynucleoside Triphosphate Triphosphohydrolase Activity of SAMHD1 Protein Contributes to the Mitochondrial DNA Depletion Associated with Genetic Deficiency of Deoxyguanosine Kinase. Journal of Biological Chemistry 290(43): 25986–25996. DOI: 10.1074/jbc.M115.675082.

Girke C, Daumann M, Niopek-Witz S, Möhlmann T (2014). Nucleobase and nucleoside transport and integration into plant metabolism. Frontiers in Plant Science 5: 443. DOI: 10.3389/fpls.2014.00443.

Goldstone DC, Ennis-Adeniran V, Hedden JJ, Groom HCT, Rice GI, Christodoulou E, et al. (2011). HIV-1 restriction factor SAMHD1 is a deoxynucleoside triphosphate triphosphohydrolase. Nature 480(7377): 379–382. DOI: 10.1038/nature10623.

Gorelova V, De Lepeleire J, Van Daele J, Pluim D, Meï C, Cuypers A, et al. (2017). Dihydrofolate reductase/thymidylate synthase fine-tunes the folate status and controls redox homeostasis in plants. The Plant Cell 29(11): 2831–2853. DOI: 10.1105/tpc.17.00433.

Govindarajan R, Leung GPH, Zhou M, Tse CM, Wang J, Unadkat JD (2009). Facilitated mitochondrial import of antiviral and anticancer nucleoside drugs by human equilibrative nucleoside transporter-3. American Journal of Physiology-Gastrointestinal and Liver Physiology 296(4): G910–G922. DOI: 10.1152/ajpgi.90672.2008.

Hammargren J, Sundström J, Johansson M, Bergman P, Knorpp C (2007). On the phylogeny, expression and targeting of plant nucleoside diphosphate kinases. Physiologia Plantarum 129(1): 79–89. DOI: 10.1111/j.1399-3054.2006.00794.x.

Heinemann KJ, Yang SY, Straube H, Medina-Escobar N, Varbanova-Herde M, Herde M, et al. (2021). Initiation of cytosolic plant purine nucleotide catabolism involves a monospecific xanthosine monophosphate phosphatase. Nature Communications 12(1): 6846. DOI: 10.1038/s41467-021-27152-4.

Herberich E, Sikorski J, Hothorn T (2010). A robust procedure for comparing multiple means under heteroscedasticity in unbalanced designs. PLoS ONE 5(3): e9788. DOI: 10.1371/journal.pone.0009788.

Hickl D, Scheuring D, Möhlmann T (2021). CTP Synthase 2 From Arabidopsis thaliana Is Required for Complete Embryo Development. Frontiers in Plant Science 12: 652434. DOI: 10.3389/fpls.2021.652434.

Hofer A, Crona M, Logan DT, Sjöberg BM (2012). DNA building blocks: keeping control of manufacture. Critical Reviews in Biochemistry and Molecular Biology 47(1): 50–63. DOI: 10.3109/10409238.2011.630372.

Hothorn T, Bretz F, Westfall P (2008). Simultaneous Inference in General Parametric Models. Biometrical Journal 50(3): 346–363. DOI: 10.1002/bimj.200810425.

Jaehme M, Slotboom DJ (2015). Structure, function, evolution, and application of bacterial Pnu-type vitamin transporters. Biological Chemistry 396(9–10): 955–966. DOI: 10.1515/hsz-2015-0113.

Kurasaka C, Nishizawa N, Ogino Y, Sato A (2022). Trapping of 5-Fluorodeoxyuridine Monophosphate by Thymidylate Synthase Confers Resistance to 5-Fluorouracil. ACS Omega 7(7): 6046–6052. DOI: 10.1021/acsomega.1c06394.

Lahouassa H, Daddacha W, Hofmann H, Ayinde D, Logue EC, Dragin L, et al. (2012). SAMHD1 restricts the replication of human immunodeficiency virus type 1 by depleting the intracellular pool of deoxynucleoside triphosphates. Nature Immunology 13(3): 223–228. DOI: 10.1038/ni.2236.

Larsen NB, Munch-Petersen B, Piškur J (2014). Tomato thymidine kinase is subject to inefficient TTP feedback regulation. Nucleosides, Nucleotides & Nucleic Acids 33(4–6): 287–290. DOI: 10.1080/15257770.2013.853781.

Law SR, Narsai R, Taylor NL, Delannoy E, Carrie C, Giraud E, et al. (2012). Nucleotide and RNA Metabolism Prime Translational Initiation in the Earliest Events of Mitochondrial Biogenesis during Arabidopsis Germination. Plant Physiology 158(4): 1610–1627. DOI: 10.1104/pp.111.192351.

Le Ret M, Belcher S, Graindorge S, Wallet C, Koechler S, Erhardt M, et al. (2018). Efficient Replication of the Plastid Genome Requires an Organellar Thymidine Kinase. Plant Physiology 178(4): 1643–1656. DOI: 10.1104/pp.18.00976.

Li H, Li L, Wu W, Wang F, Zhou F, Lin Y (2022). SvSTL1 in the large subunit family of ribonucleotide reductases plays a major role in chloroplast development of Setaria viridis. The Plant Journal 111(3): 625–641. DOI: 10.1111/tpj.15842.

Liu B, Czajka A, Malik AN, Hussain K, Jones PM, Persaud SJ (2015). Equilibrative nucleoside transporter 3 depletion in β-cells impairs mitochondrial function and promotes apoptosis: Relationship to pigmented hypertrichotic dermatosis with insulin-dependent diabetes. Biochimica et Biophysica Acta - Molecular Basis of Disease 1852(10 Pt A): 2086–2095. DOI: 10.1016/j.bbadis.2015.07.002.

Livak KJ, Schmittgen TD (2001). Analysis of Relative Gene Expression Data Using Real-Time Quantitative PCR and the 2−ΔΔCT Method. Methods 25(4): 402–408. DOI: 10.1006/meth.2001.1262.

Lu C, Wang Q, Jiang Y, Zhang M, Meng X, Li Y, et al. (2023). Discovery of a novel nucleoside immune signaling molecule 2′-deoxyguanosine in microbes and plants. Journal of Advanced Research 46: 1–15. DOI: 10.1016/j.jare.2022.06.014.

MacVicar T (2025). High tide or low tide: the transport and metabolism of mitochondrial nucleotides. Biochemical Journal 482(16): 1105–1122. DOI: 10.1042/BCJ20253237.

Marx A, Alian A (2015). The first crystal structure of a dTTP-bound deoxycytidylate deaminase validates and details the allosteric-inhibitor binding site. Journal of Biological Chemistry 290(1): 682–690. DOI: 10.1074/jbc.M114.617720.

Møller IM, Rao RSP, Jiang Y, Thelen JJ, Xu D (2020). Proteomic and Bioinformatic Profiling of Transporters in Higher Plant Mitochondria. Biomolecules 10(8): 1190. DOI: 10.3390/biom10081190.

Mori R, Ukai J, Tokumaru Y, Niwa Y, Futamura M (2022). The mechanism underlying resistance to 5-fluorouracil and its reversal by the inhibition of thymidine phosphorylase in breast cancer cells. Oncology Letters 24(3): 311. DOI: 10.3892/ol.2022.13431.

Morley SA, Ahmad N, Nielsen BL (2019). Plant Organelle Genome Replication. Plants 8(10): 358. DOI: 10.3390/plants8100358.

Nájera-Martínez M, Pedroza-García JA, Suzuri-Hernández LJ, Mazubert C, Drouin-Wahbi J, Vázquez-Ramos J, et al. (2020). Maize Thymidine Kinase Activity Is Present throughout Plant Development and Its Heterologous Expression Confers Tolerance to an Organellar DNA-Damaging Agent. Plants 9(8): 930. DOI: 10.3390/plants9080930.

Niehaus M, Straube H, Specht A, Baccolini C, Witte CP, Herde M (2022). The nucleotide metabolome of germinating Arabidopsis thaliana seeds reveals a central role for thymidine phosphorylation in chloroplast development. The Plant Cell 34(10): 3790–3813. DOI: 10.1093/plcell/koac207.

Niu M, Wang Y, Wang C, Lyu J, Wang Y, Dong H, et al. (2017). ALR encoding dCMP deaminase is critical for DNA damage repair, cell cycle progression and plant development in rice. Journal of Experimental Botany 68(21–22): 5773–5786. DOI: 10.1093/jxb/erx380.

Nordlund P, Reichard P (2006). Ribonucleotide reductases. Annual Review of Biochemistry 75: 681–706. DOI: 10.1146/annurev.biochem.75.103004.142443.

Ohler L, Niopek-Witz S, Mainguet SE, Möhlmann T (2019). Pyrimidine Salvage: Physiological Functions and Interaction with Chloroplast Biogenesis. Plant Physiology 180(4): 1816– 1828. DOI: 10.1104/pp.19.00329.

Pedroza-García JA, Nájera-Martínez M, Sánchez MP, Plasencia J (2015). Arabidopsis thaliana thymidine kinase 1a is ubiquitously expressed during development and contributes to confer tolerance to genotoxic stress. Plant Molecular Biology 87(3): 303–315. DOI: 10.1007/s11103-014-0277-7.

Pedroza-García JA, Nájera-Martínez M, Mazubert C, Aguilera-Alvarado P, Drouin-Wahbi J, Sánchez-Nieto S, et al. (2019). Role of pyrimidine salvage pathway in the maintenance of organellar and nuclear genome integrity. The Plant Journal 97(3): 430–446. DOI: 10.1111/tpj.14128.

Rinne J, Witte CP, Herde M (2021). Loss of MAR1 Function is a Marker for Co-Selection of CRISPR-Induced Mutations in Plants. Frontiers in Genome Editing 3: 723384. DOI: 10.3389/fgeed.2021.723384.

Rinne J, Niehaus M, Medina-Escobar N, Straube H, Schaarschmidt F, Rugen N, et al. (2024). Three Arabidopsis UMP kinases have different roles in pyrimidine nucleotide biosynthesis and (deoxy)CMP salvage. The Plant Cell 36(9): 3611–3630. DOI: 10.1093/plcell/koae170.

Roa H, Lang J, Culligan KM, Keller M, Holec S, Cognat V, et al. (2009). Ribonucleotide reductase regulation in response to genotoxic stress in Arabidopsis. Plant Physiology 151(1): 461– 471. DOI: 10.1104/pp.109.140053.

Ronceret A, Gadea-Vacas J, Guilleminot J, Lincker F, Delorme V, Lahmy S, et al. (2008). The first zygotic division in Arabidopsis requires de novo transcription of thymidylate kinase. The Plant Journal 53(5): 776–789. DOI: 10.1111/j.1365-313X.2007.03372.x.

Sarmiento-Mañús R, Fontcuberta-Cervera S, González-Bayón R, Hannah MA, Álvarez-Martínez FJ, Barrajón-Catalán E, et al. (2023). Analysis of the Arabidopsis venosa4-0 mutant supports the role of VENOSA4 in dNTP metabolism. Plant Science 335: 111819. DOI: 10.1016/j.plantsci.2023.111819.

Schindelin J, Arganda-Carreras I, Frise E, Kaynig V, Longair M, Pietzsch T, et al. (2012). Fiji: an open-source platform for biological-image analysis. Nature Methods 9(7): 676–682. DOI: 10.1038/nmeth.2019.

Straube H, Niehaus M, Zwittian S, Witte CP, Herde M (2021). Enhanced nucleotide analysis enables the quantification of deoxynucleotides in plants and algae revealing connections between nucleoside and deoxynucleoside metabolism. The Plant Cell 33(2): 270–289. DOI:10.1093/plcell/koaa028.

Straube H, Straube J, Rinne J, Fischer L, Niehaus M, Witte CP, et al. (2023). An inosine triphosphate pyrophosphatase safeguards plant nucleic acids from aberrant purine nucleotides. New Phytologist 237(5): 1759–1775. DOI: 10.1111/nph.18656.

Sweetlove LJ, Mowday B, Hebestreit HF, Leaver CJ, Millar AH (2001). Nucleoside diphosphate kinase III is localized to the inter-membrane space in plant mitochondria. FEBS Letters 508(2): 272–276. DOI: 10.1016/S0014-5793(01)03069-1.

Traba J, Satrústegui J, del Arco A (2011). Adenine nucleotide transporters in organelles: novel genes and functions. Cellular and Molecular Life Sciences 68(7): 1183–1206. DOI: 10.1007/s00018-010-0612-3.

Villarroya J, Lara MC, Dorado B, Garrido M, García-Arumí E, Meseguer A, et al. (2011). Targeted impairment of thymidine kinase 2 expression in cells induces mitochondrial DNA depletion and reveals molecular mechanisms of compensation of mitochondrial respiratory activity. Biochemical and Biophysical Research Communications 407(2): 333–338. DOI: 10.1016/j.bbrc.2011.03.018.

Wang C, Liu Z (2006). Arabidopsis ribonucleotide reductases are critical for cell cycle progression, DNA damage repair, and plant development. The Plant Cell 18(2): 350–365. DOI: 10.1105/tpc.105.037044.

Wang H, Tu R, Ruan Z, Wu D, Peng Z, Zhou X, et al. (2022). STRIPE3, encoding a human dNTPase SAMHD1 homolog, regulates chloroplast development in rice. Plant Science 323: 111395. DOI: 10.1016/j.plantsci.2022.111395.

Welin M, Kosinska U, Mikkelsen NE, Carnrot C, Zhu C, Wang L, et al. (2004). Structures of thymidine kinase 1 of human and mycoplasmic origin. Proceedings of the National Academy of Sciences of the United States of America 101(52): 17970–17975. DOI: 10.1073/pnas.0406332102.

Witte CP, Herde M (2020). Nucleotide Metabolism in Plants. Plant Physiology 182(1): 63–78. DOI: 10.1104/pp.19.00955.

Xu J, Deng Y, Li Q, Zhu X, He Z (2014). STRIPE2 Encodes a Putative dCMP Deaminase that Plays an Important Role in Chloroplast Development in Rice. Journal of Genetics and Genomics 41(10): 539–548. DOI: 10.1016/j.jgg.2014.05.008.

Xu D, Leister D, Kleine T (2020). VENOSA4, a Human dNTPase SAMHD1 Homolog, Contributes to Chloroplast Development and Abiotic Stress Tolerance. Plant Physiology 182(2): 721– 729. DOI: 10.1104/pp.19.01108.

Yoshida Y, Sarmiento-Mañús R, Yamori W, Ponce MR, Micol JL, Tsukaya H (2018). The Arabidopsis phyB-9 Mutant Has a Second-Site Mutation in the VENOSA4 Gene That Alters Chloroplast Size, Photosynthetic Traits, and Leaf Growth. Plant Physiology 178(1): 3–6. DOI: 10.1104/pp.18.00764.

Zeileis A (2004). Econometric Computing with HC and HAC Covariance Matrix Estimators. Journal of Statistical Software 11(10): 1–17. DOI: 10.18637/jss.v011.i10.

Zeileis A, Köll S, Graham N (2020). Various Versatile Variances: An Object-Oriented Implementation of Clustered Covariances in R. Journal of Statistical Software 95(1): 1–36. DOI: 10.18637/jss.v095.i01.

