## Supplementary Figures and Tables for "Deoxyribonucleotide dephosphorylation by VENOSA4 supports organellar genome replication in plants"

**Supplemental Figures and Tables**

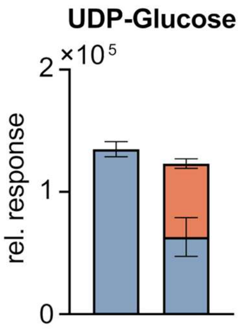


**Supplemental Figure 1. UDP-Glucose in germinating Arabidopsis seeds supplied with $^{15}N$-uridine.**

Col-0 seeds were grown on ½ MS medium with or without $^{15}N$-uridine and harvested after 48 h. Stacked bars show relative response of UDP-Glucose in the same samples as in Fig. 1: unlabeled (blue, lower column part) and $^{15}N$-labeled metabolite species (orange, upper column part). For each metabolite, the left bar represents control conditions (no $^{15}N$-uridine) and the right bar treatment with $^{15}N$-uridine. Data are means ± SD (*n* = 4).

**Supplemental Table 1. Percentages of unlabeled and labeled metabolite species upon $^{15}N$-uridine feeding.**

Col-0 seeds were grown on ½ MS medium supplemented with $^{15}N$-uridine and harvested after 48 h. The relative proportions of unlabeled and $^{15}N$-labeled metabolite species were calculated for individual replicates relative to the total metabolite amount (unlabeled + labeled = 100%) and averaged across replicates.

| Metabolite | Unlabeled (%) | Labeled (%) |
|---|---|---|
| CDP | 57,1% | 42,9% |
| CMP | 68,9% | 31,1% |
| CTP | 53,6% | 46,4% |
| dT | 63,7% | 36,3% |
| dTDP | 43,6% | 56,4% |
| dTMP | 43,5% | 56,5% |
| dTTP | 49,9% | 50,1% |
| UDP-Glucose | 50,6% | 49,4% |

| | | |
|---|---|---|
| UDP | 43,9% | 56,1% |
| UMP | 45,7% | 54,3% |
| Uridine | 2,4% | 97,6% |
| UTP | 46,0% | 54,0% |

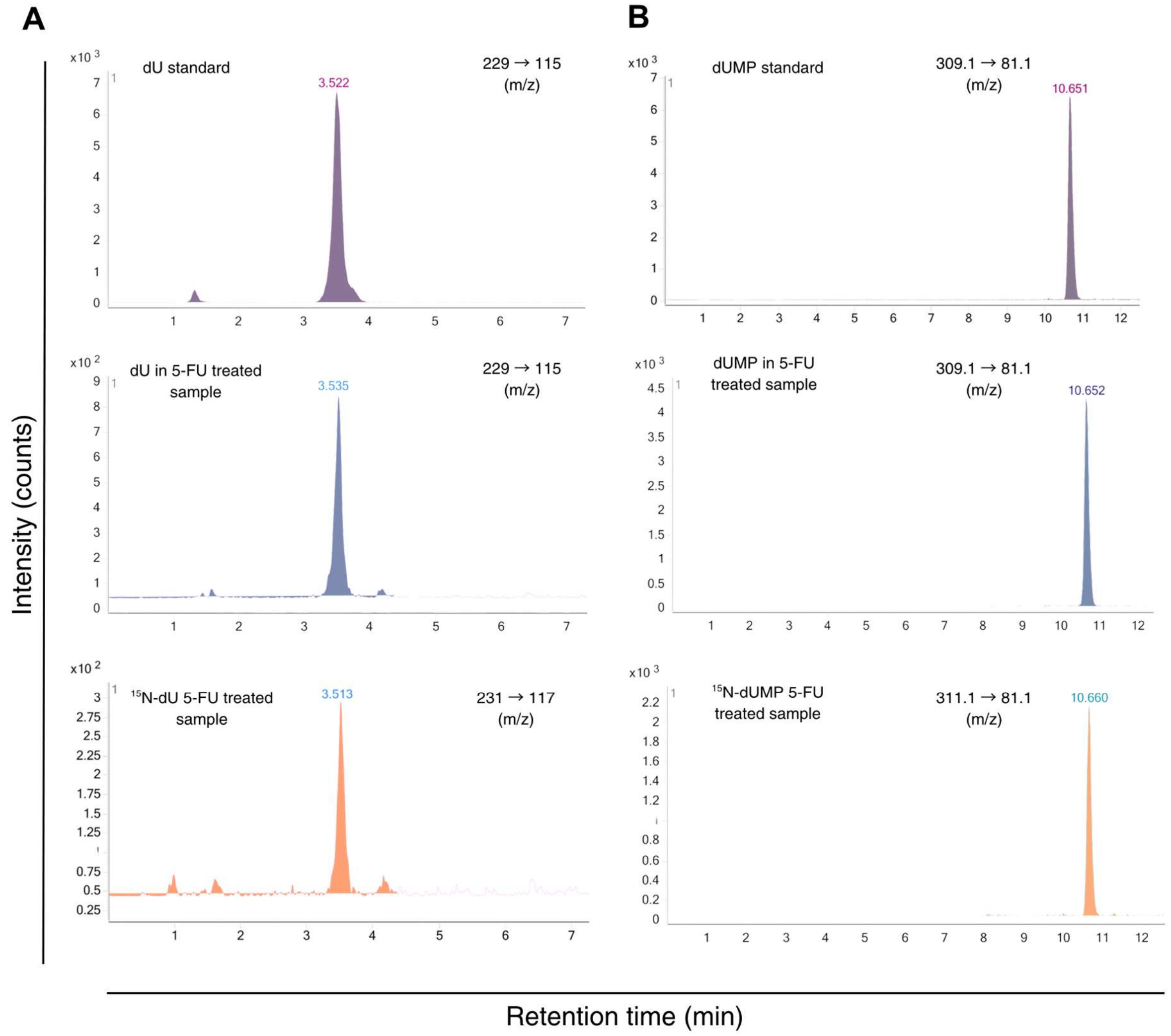


**Supplemental Figure 2. Chromatograms of dU and dUMP peaks of the standard and in plant matrix.**

**(A)** Representative multiple-reaction monitoring (MRM) chromatograms of dU standard (top panel), dU (middle panel) and $^{15}$N-dU in matrix of *A. thaliana* seedlings. Retention time of the standard (3.522 min) matches those of the dU and $^{15}$N-dU peaks.

**(B)** Representative (MRM) chromatograms of dUMP standard (top panel), dUMP in plant matrix (middle panel) and $^{15}N$-dUMP in plant matrix. Retention time of the standard (10.651 min) matches those of the dUMP and $^{15}N$-dUMP peaks.

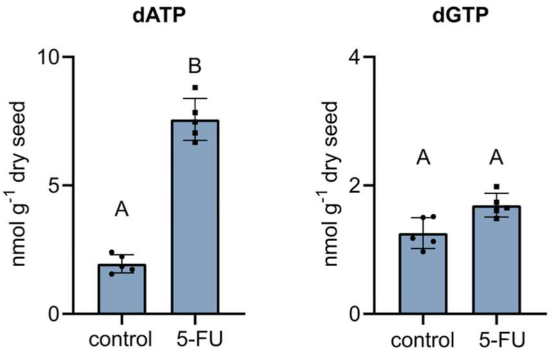


**Supplemental Figure 3. Effects of 5-FU on dGTP and dATP in germinating Arabidopsis seeds.** Col-0 seeds were grown on medium containing either 0 or 50 µM 5-fluorouridine (5-FU) and harvested after 48 h after transfer to grwoth conditions. dATP and dGTP were quantified in unlabeled control and unlabeled 5-FU treated samples of Fig. 4. Data are means ± SD (*n* = 5); different letters indicate statistically significant differences between treatments ($P < 0.05$).

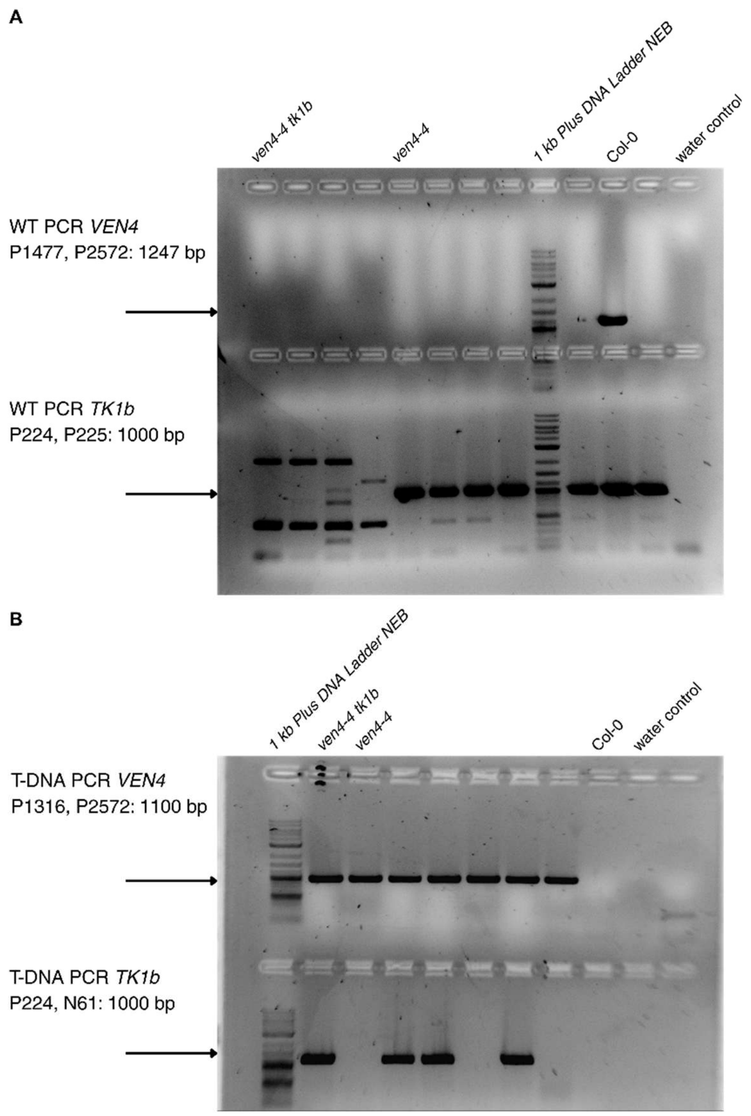


**Supplemental Figure 4. Genotyping of *ven4-4* mutant and *ven4-4 tk1b* double-mutant plants.**

Agarose gels showing PCR genotyping of the *VEN4* and *TK1b* loci. (A) Wild-type allele-specific PCR reactions for *VEN4* (upper gel part) and *TK1b* (lower gel part). (B) T-DNA insertion-specific PCR reactions for *VEN4* (upper gel part) and *TK1b* (lower gel part).

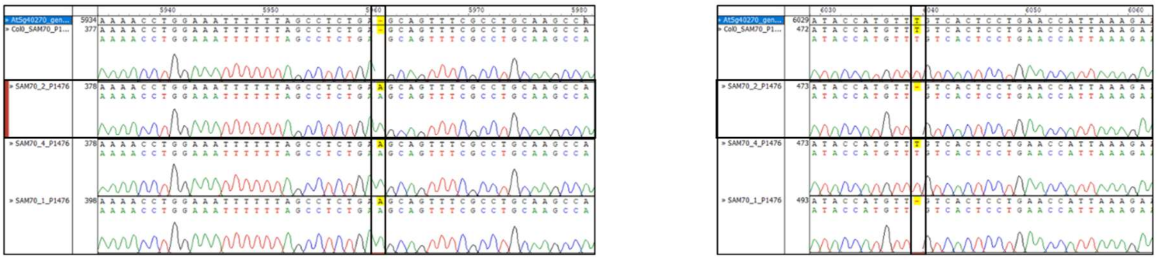

**Supplemental Figure 5. Sequence validation of the CRISPR/Cas9-generated *ven4-5* mutant.** Sequencing chromatograms of the *VEN4* (*At5g40270*) locus at the CRISPR/Cas9 target sites. Sequences of Col-0 and CRISPR-edited plants were aligned to the *VEN4* reference sequence. Sequence alterations at the targeted positions are highlighted in yellow.

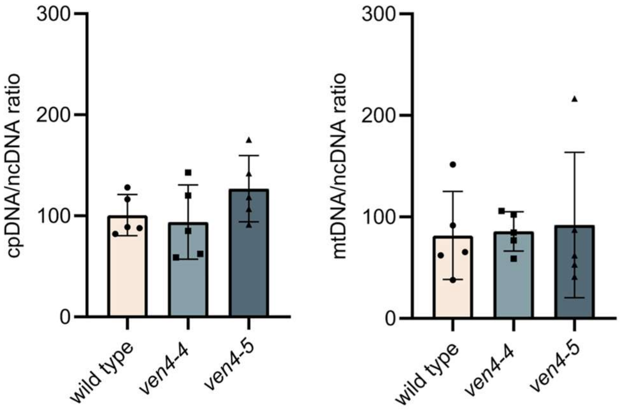


**Supplemental Figure 6. Organelle DNA abundance in dry seeds of *VEN4* mutants.**
Organellar to nuclear DNA ratios (cpDNA/nDNA and mtDNA/nDNA) in dry seeds of wild type, *ven4-4* and *ven4-5* plants. Organelle and nuclear DNA abundance were quantified by qPCR using primer pairs targeting the plastid gene RBCL (cpDNA), the mitochondrial gene COX1 (mtDNA) and the nuclear gene UBC21 (ncDNA). cpDNA/ncDNA and mtDNA/ncDNA ratios were derived from the corresponding Ct values. Data are means ± SD (*n* = 5); different letters indicate statistically significant differences between genotypes (P < 0.05).

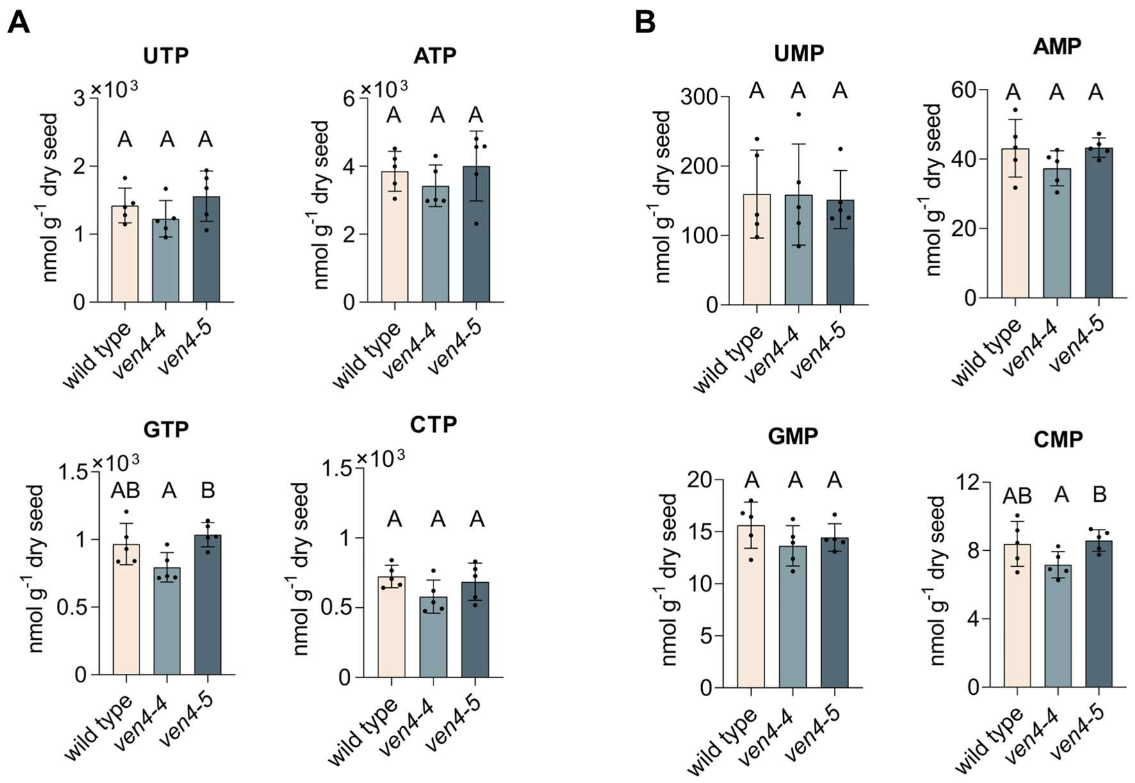


**Supplemental Figure 7. rNTP and rNMP concentrations in *VEN4* mutants during GSE.**

UTP, ATP, GTP, CTP and UMP, AMP, GMP, CMP concentrations in 48-hour-old wild type, *ven4-4* and *ven4-5* plants grown on liquid ½ MS medium. Data are means ± SD ($n$ = 5); different letters indicate statistically significant differences between genotypes ($P < 0.05$).

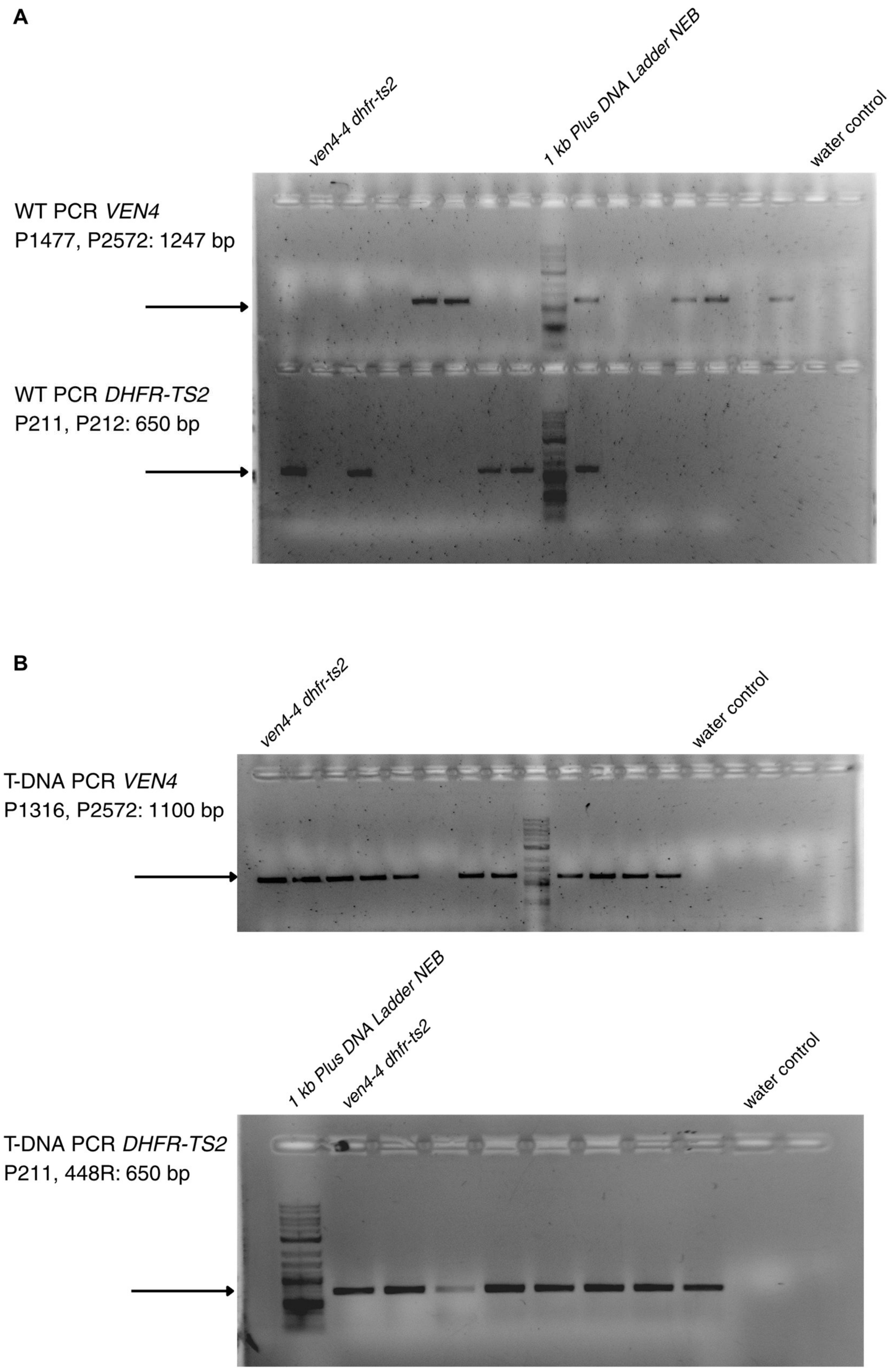


**Supplemental Figure 8. Genotyping of *ven4-4 dhfr-ts2* double-mutant plants.**

Agarose gels showing PCR genotyping of the *VEN4* and *DHFR-TS2* loci. (A) Wild-type allele-specific PCR reactions for *VEN4* (upper gel part) and *DHFR-TS2* (lower gel part). (B) T-DNA insertion-specific PCR reactions for *VEN4* (upper gel) and *DHFR-TS2* (lower gel).

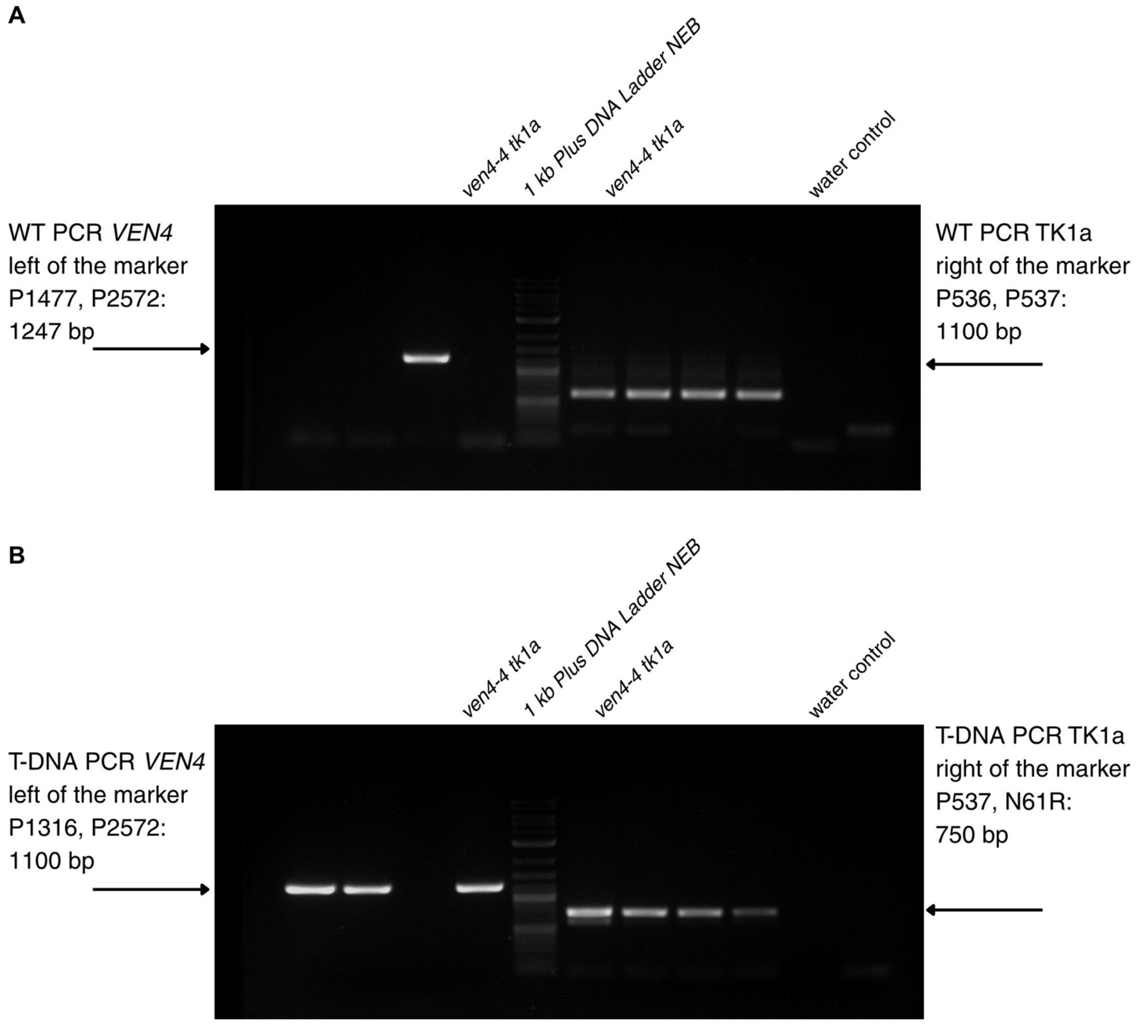


**Supplemental Figure 9. Genotyping of *ven4-4 tk1a* double-mutant plants.**

Agarose gels showing PCR genotyping of the *VEN4* and *TK1a* loci. (A) Wild-type allele-specific PCR reactions for *VEN4* (left of the marker) and *TK1a* (right of the marker). (B) T-DNA insertion-specific PCR reactions for *VEN4* (left of the marker) and *TK1a* (right of the marker).
